# Expanding the substrate selectivity of the fimsbactin biosynthetic adenylation domain, FbsH

**DOI:** 10.1101/2024.07.26.605314

**Authors:** Syed Fardin Ahmed, Adam Balutowski, Jinping Yang, Timothy A. Wencewicz, Andrew M. Gulick

## Abstract

Nonribosomal peptide synthetases (NRPSs) produce diverse natural products including siderophores, chelating agents that many pathogenic bacteria produce to survive in low iron conditions. Engineering NRPSs to produce diverse siderophore analogs could lead to the generation of novel antibiotics and imaging agents that take advantage of this unique iron uptake system in bacteria. The highly pathogenic and antibiotic-resistant bacteria *Acinetobacter baumannii* produces fimsbactin, an unusual branched siderophore with iron-binding catechol groups bound to a serine or threonine side chain. To explore the substrate promiscuity of the assembly line enzymes, we report a structure-guided investigation of the stand-alone aryl adenylation enzyme FbsH. We report on structures bound to its native substrate 2,3-dihydroxybenzoic acid (DHB) as well as an inhibitor that mimics the adenylate intermediate. We produced enzyme variants with an expanded binding pocket that are more tolerant for analogs containing a DHB C4 modification. Wild-type and mutant enzymes were then used in an *in vitro* reconstitution analysis to assess the production of analogs of the final product as well as several early-stage intermediates. This analysis shows that some altered substrates progress down the fimsbactin assembly line to the downstream domains. However, analogs from alternate building blocks are produced at lower levels, indicating that selectivity exists in the downstream catalytic domains. These findings expand the substrate scope of producing condensation products between serine and aryl acids and identify the bottlenecks for chemoenzymatic production of fimsbactin analogs.

## Introduction

Siderophores are iron-chelating natural products produced by microorganisms such as bacteria and fungi. These low molecular weight compounds have very high binding affinities for iron to promote the growth of bacteria in low iron environments.^1^ Different siderophores are classified by the catechol, hydroxamate, phenolate, and carboxylate groups used to coordinate iron. After binding ferric ion, the complexed siderophores are recognized by high affinity surface receptor proteins to transport the iron-loaded siderophore to enter the cell.^2^ Once inside the cell, the iron is released by the breakdown of the siderophore backbone or the reduction of the iron and becomes bioavailable for metabolic functions.^3^ Due to the importance of siderophores for the growth of bacteria in low iron environments, these small molecules are essential virulence factors for multi-drug resistant (MDR) bacterial pathogens such as *Acinetobacter baumannii*.^4, 5^ This unique bacterial iron uptake strategy has been targeted for the development of new therapeutics and diagnostics including antivirulence agents that block siderophore biosynthesis and utilization, siderophore-antibiotic conjugates (sideromycins) for targeted drug delivery, and siderophores labeled with a molecular tracer for infection imaging.^4-10^

Bacterial siderophores are primarily produced by nonribosomal peptide synthetases (NRPSs) or NRPS-independent siderophore (NIS) synthetase pathways.^11^ NRPSs are specialized multi-domain proteins that function in an assembly-line manner to produce peptide chains in the absence of ribosomes. They consist of several key domains, each specialized to carry out a specific function in synthesizing these peptides.^12-15^ These domains include the adenylation domain (A), which activates the substrate using ATP and loads it onto the pantetheine thiol group of a peptidyl carrier protein (PCP, which is also known as the thiolation domain, T). PCP domains are covalently modified with a coenzyme A-derived pantetheine prosthetic group to which substrate building blocks are bound as a thioester. The adenylation domains selectively activate specific amino or aryl acids.^16^ Once loaded onto the PCP domains, the PCP domain delivers the activated substrate to the condensation domain (C), which catalyzes peptide bond formation with a free amino group from a downstream tethered amino acid.^17^ Finally, a terminal thioesterase (TE) domain catalyzes product release via cyclization or hydrolysis.^18^ NRPS pathways may include other specialized domains such as epimerase,^19^ methyltransferase,^20^ and cyclization domains that play roles in the condensation, cyclization, and dehydration of serine, threonine, and cysteine residues.^17, 21^ Conventional NRPS assembly lines are classified into modules, with each module playing a role in synthesizing one peptide bond in the growing peptidyl chain. While such linear domain and modular architecture allows the prediction of NRPS products based on the protein sequences alone, the existence of nonlinear NRPS pathways poses challenges to the structural prediction of the nonribosomal peptide products.

Due to their highly selective nature, NRPS adenylation domains, including aryl adenylating enzymes, have been of interest for both structural and functional characterization.^16, 22^ The adenylation domains contain two subdomains, an N-terminal A_core_ domain that encompasses the first ∼400-450 residues and a smaller ∼120 residue C-terminal A_sub_ domain. Adenylation domains catalyze the substrate activation and loading with a two-step ping-pong reaction mechanism. In the adenylate-forming step, the aryl-adenylation domain recognizes an aryl acid and activates it using ATP, forming an aryl adenylate and releasing pyrophosphate. In each adenylation domain, substrate recognition is governed by a group of residues that line the binding pocket, which can be deciphered as substrate specificity codes.^23, 24^ Once the aryl-adenylate intermediate is produced, a second thioester-forming step transfers the aryl acid to the phosphopantetheine arm of the corresponding PCP domain.^25^ To carry out this two-step reaction, the A_sub_ domain rotates by ∼140° to adopt two unique conformations, namely the adenylate-forming conformation and a thioester-forming conformation.^22, 26^ Representative structures of stand-alone aryl adenylating enzymes have been solved where the A_sub_ is present in the adenylation conformation,^27, 28^ the thioester conformation in complex with the partner PCP domain,^29-32^ and multiple intermediate conformations;^33^ additionally, in the other structures, this dynamic subdomain is disordered and likely adopts multiple orientations within the crystal lattice.^34^

In addition to exploring the reaction mechanism and domain conformations, studies e of substrate recognition and selectivity of aryl-adenylation domains have shown significant conservation of binding pocket residues in aryl acid binding adenylation domains found across different catechol siderophore biosynthetic pathways.^27, 30, 33-38^ Structures of adenylation domains bound to non-native substrates also support efforts to understand the determinants of substrate promiscuity and to design novel inhibitors.^28, 34^ The substrate-recognizing residues in the binding pocket of the stand-alone adenylation domain EntE from *E. coli* show that Asn235 and Ser240 form hydrogen bonds with the two hydroxyl groups of DHB, whereas Tyr236, Val331, and Val339 form the base of a hydrophobic pocket for DHB.^35^

Efforts to expand the promiscuity for non-native substrates can potentially allow a chemoenzymatic approach to produce siderophore derivatives with therapeutic potentials.^39^ As NRPS adenylation domains serve as “gatekeepers” for the selection of starter substrates, they have been attractive targets for engineering for the development of novel natural products with therapeutic potential.^40^ These studies integrate structural information to enhance the rational engineering of adenylation domains that can incorporate substrate analogs to generate new products. Point mutations in EntE from enterobactin biosynthesis have expanded the binding pocket to tolerate a diverse range of benzoic acid analogs.^35^ Further site-directed mutagenesis was utilized to enhance EntE for greater activity for a non-native substrate salicylic acid, over its native substrate DHB.^36^

Here, we expand these studies by investigating the biosynthesis of fimsbactin, a siderophore produced by *Acinetobacter baumannii*.^41^ Previously, fimsbactin and synthetic analogs have been shown to outcompete the essential siderophore acinetobactin for uptake in *A. baumannii* providing a route to siderophore-based antivirulence agents.^42, 43^ Fimsbactin is assembled by four NRPS enzymes encoded by the genes *fbsEFGH* (Figure 1).^41^ Previous studies have explored the fimsbactin biosynthetic pathway and proposed an unusual branching mechanism for the formation of fimsbactin from the precursor molecules DHB, L-Ser, and *N*-acetyl-*N*-hydroxyputrescine (ahPut).^44^ The fimsbactin pathway involves a freestanding adenylation domain FbsH as well as three multidomain proteins FbsE, FbsF, and FbsG, each of which contains one or two carrier protein domains. The N-terminal PCP of FbsE is predicted to be the partner for FbsH, enabling the loading of the DHB onto this aryl carrier protein (ArCP) domain. As FbsE contains a truncated adenylation domain, the serine substrates are predicted to be incorporated *in trans* by the activity of the FbsF adenylation domain to be loaded onto FbsE C-terminal PCP and FbsG PCP domains. The tandem cyclization domains and condensation domain are responsible for the production of the oxazoline, the *O-*DHB-Ser, and the final release of full-length fimsbactin A, although the exact time and order of reactions remains unclear.

**Figure 1:**
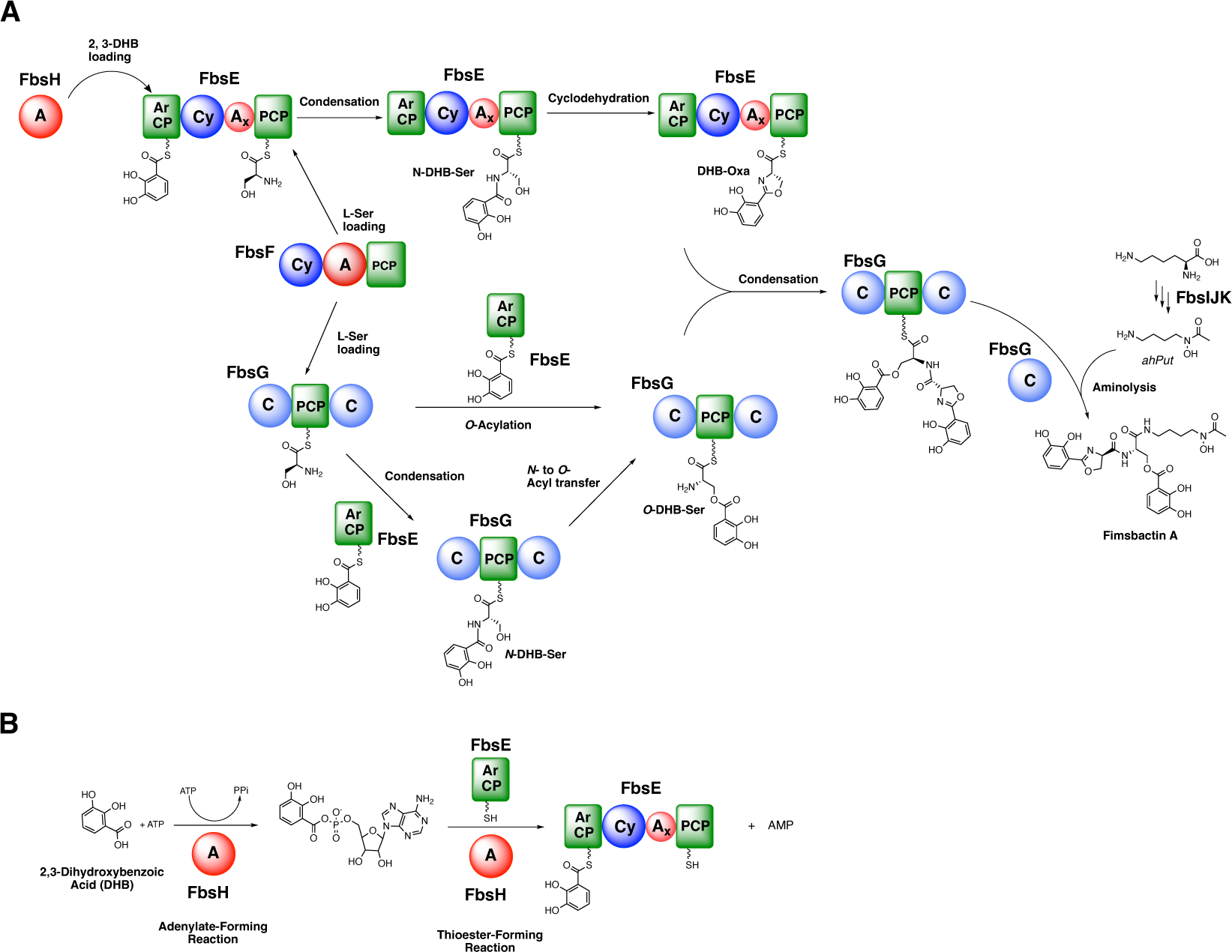
Biosynthesis of Fimsbactin. (A) Proposed mechanism of fimsbactin biosynthesis by NRPSs FbsEFGHIJK. (B) Two-step adenylation reaction catalyzed by a standalone adenylation domain FbsH.

Here, we present two new structures of the aryl adenylation domain FbsH bound to its native substrate DHB and the inhibitor 5’-*O*-[*N*-(salicyl)sulfamoyl]adenosine (Sal-AMS).^45^ We identified several binding pocket residues of FbsH and mutated them to expand its pocket size. Using substrate screening and steady-state kinetic analysis, we demonstrated that an expanded binding pocket of FbsH can tolerate several larger DHB analogs and show significant improvement in activity in adenylation assays compared to wild-type FbsH. Finally, by implementing full *in vitro* reconstitution experiments, we show that we can incorporate certain DHB analogs through the biosynthetic pathway to generate fimsbactin analogs harboring two alternate aryl groups. While this demonstrates the ability of the downstream domains to also accommodate the new building blocks, the selection of intermediates in the subsequent condensation and cyclization steps limits the production of fimsbactin analogs compared to their natural counterpart.

## Results

### Structures of FbsH bound to DHB and Salicyl-AMS

To better understand the substrate selectivity of FbsH, we solved the structures of FbsH bound to its native substrate DHB and to the inhibitor Sal-AMS. Both structures were solved in the *P*1 space group, each having two molecules per asymmetric unit. The Sal-AMS-bound structure was first solved using an AlphaFold model^46^ of the N terminal subdomain, A_core_, as the molecular replacement search model. Upon preliminary inspection, it was observed that the dynamic C-terminal subdomain had a weaker density than the A_core_ subdomain. The A_sub_ domains were manually built for both chains and observed to be in an intermediate conformation in between the two catalytic conformations (Figure 2A, B). In addition, the superposition of each chain shows that the A_sub_ domain of chain A was in a slightly different conformation than that of Chain B. A model of the protein atoms from the A_core_ subdomain of the Sal-AMS-bound structure was then used to solve the DHB-bound structure. Strong density was observed at the active site binding pockets of both structures (Supplemental Figure 1), corresponding to the respective ligands. The DHB co-crystal structure was determined from a crystal that also contained AMP. In chain A, weak density was present in the nucleotide binding pocket that may represent low occupancy of AMP; this density was left unmodeled in the final model.

**Figure 2:**
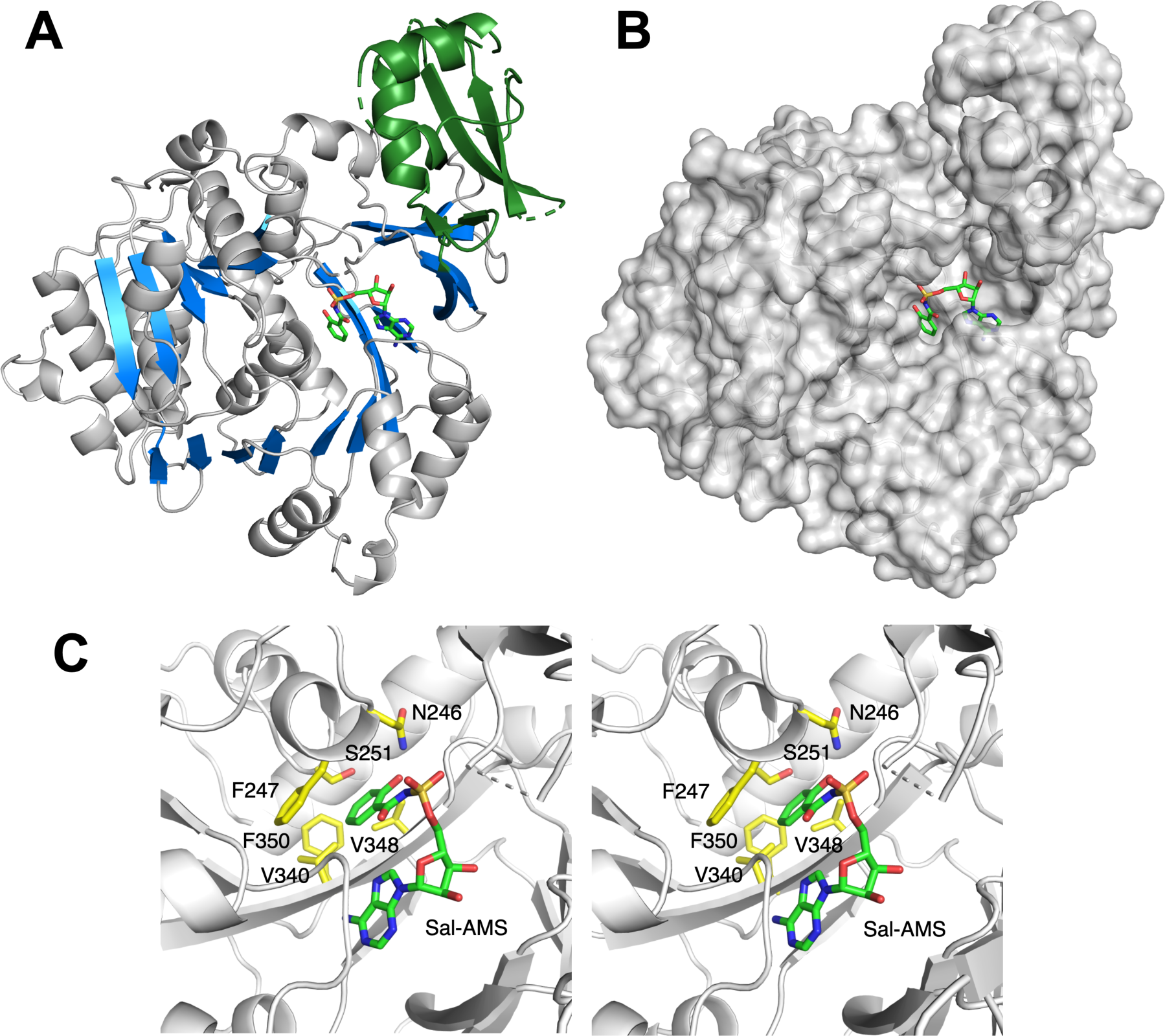
Structure of FbsH bound to Sal-AMS. (A) Overall structure of FbsH bound to Sal-AMS. Dynamic C-term subdomain (shown in green) is present in an intermediate conformation. (B) Surface representation of FbsH shows that the binding pocket forms a hollow cavity for substrates to bind. (C) Stereorepresantation of the substrate binding pocket of FbsH shows conserved residues across aryl-adenylating enzymes. N246 and S251 form hydrogen bonds with the hydroxyl groups of catechol substrates. These bonds are necessary to bind the substrate. F247, V340, V348 and F350 form the base of the pocket and form hydrophobic interactions with the substrate.

FbsH was observed to have structural homology with previously solved structures of aryl acid binding adenylation domains (Supplemental Table 1). Compared to the N-terminal A_core_ domain of EntE (PDB **3RG2**),^30^ FbsH has a root-mean-square displacement of 0.79 Å over 438 residues and a sequence similarity of 39.6%. In addition, the binding pocket residues Asn246 and Ser251 were conserved in hydrogen bonding distances from the hydroxyl groups of the bound DHB. These residues are conserved across different catechol-binding A-domains, including BasE,^34^ and are necessary for the recognition and binding of substrates. Finally, other binding pocket residues are conserved with previously solved structures, including Phe247, Val340, Val348, Asn349, and Phe350. These residues form hydrophobic interactions with the bound substrates (Supplemental Figure 2).

### Enzymatic Activity of FbsH toward DHB analogs

To better understand the substrate selectivity of FbsH towards DHB analogs, we explored the catalytic activity of wild-type FbsH with these analogs. We first screened for substrate selectivity based on measuring the specific activity of FbsH in the presence of a variety of substrates (Figure 3A). In the specific activity screen, FbsH showed similar activities for both DHB and salicylic acid, demonstrating that the removal of the hydroxyl from the C3 position has a limited impact on substrate adenylation. This prompted us to explore activity with several analogs substituted at the C3- and C4-positions. Initial screening revealed an overall preference of wild-type FbsH towards C3-substituted analogs over C4-substituted analogs presumably due to a lack of space at the base of the DHB-binding pocket. In fact, no activity was observed with several 4-substituted analogs such as 4-nitro and 4-azido salicylic acid.

**Figure 3:**
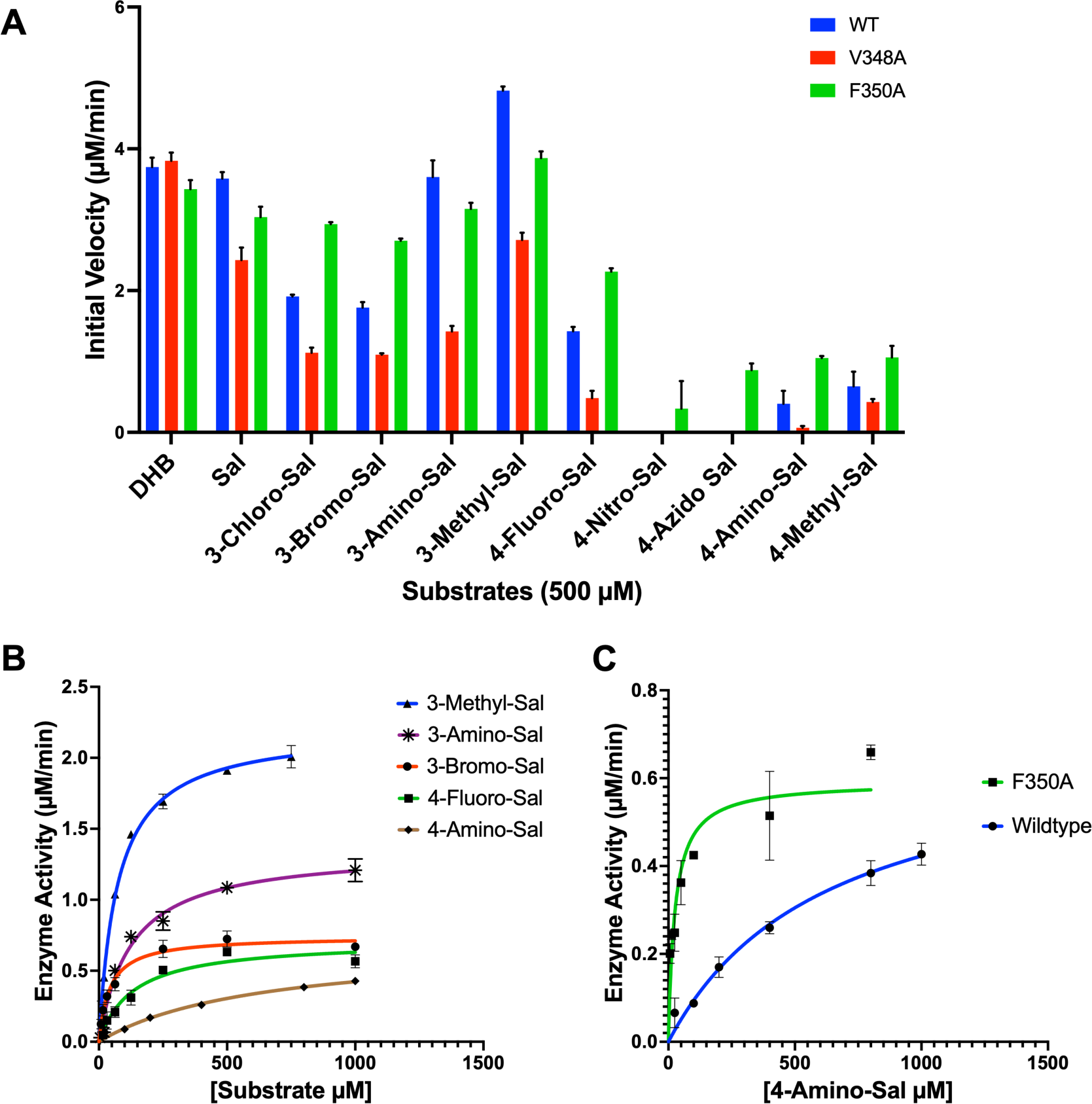
Specificity Screening and steady-state kinetic analysis of wild-type and mutant FbsH with DHB analogs. (A) Specific activity screening of wild-type FbsH showed an overall preference for salicylic acid and C3-substituted DHB analogs over C4-substituted analogs. (B) Steady-state kinetic analysis with wild-type FbsH. (C) The F350A mutant that expands the binding pocket of FbsH, significantly improved the overall catalytic efficiencies with the 4-substituted analogs, with a ∼53-fold improvement for 4-Amino-Sal.

We measured steady-state kinetic data to explore the binding affinity and catalytic efficiency of FbsH with a selection of analogs (Figure 3B). Upon comparing apparent kinetic constants (Table 1, Supplemental Figure 3), we observed that the C3-substituted analogs had catalytic efficiencies 3- to 5-fold lower, compared to DHB, driven by both reduced catalytic turnover (apparent k_cat_) and binding affinities (apparent K_M_), with values ranging from 27–140 μM for the C3-substituted analogs in comparison to 21 ± 3 μM for DHB. With the C4-substituted analogs, a more significant decline in catalytic efficiency was observed. This includes more than a 250-fold decline in catalytic efficiency for 4-amino salicylic acid and over a 100-fold decline for 4-methyl salicylic acid when compared to DHB, as indicated by both an increase in apparent K_M_ values as well as a decrease in apparent k_cat_.

**Table 1:** Apparent steady-state kinetics of wild-type and mutant enzyme with different substrates.

| Protein | Substrate | Protein Conc. ($\mu M$) | Vmax ($\mu M/min$) | $K_M$ ($\mu M$) | $k_{cat}$ ($sec^{-1}$) | $k_{cat}/K_M$ ($M^{-1} sec^{-1}$) |
| --- | --- | --- | --- | --- | --- | --- |
| FbsH WT | DHB | 1 | $1.17 \pm 0.05$ | $20.6 \pm 2.60$ | $(1.95 \pm 0.08) \times 10^{-2}$ | $9.46 \times 10^2$ |
| FbsH WT | Sal | 1 | $1.38 \pm 0.05$ | $71.9 \pm 11.2$ | $(2.30 \pm 0.09) \times 10^{-2}$ | $3.19 \times 10^2$ |
| FbsH WT | 3-Chloro-Sal | 1 | $0.74 \pm 0.02$ | $30.7 \pm 3.36$ | $(1.23 \pm 0.03) \times 10^{-2}$ | $3.98 \times 10^2$ |
| FbsH WT | 3-Bromo-Sal | 1 | $0.74 \pm 0.02$ | $41.9 \pm 4.87$ | $(1.23 \pm 0.04) \times 10^{-2}$ | $2.94 \times 10^2$ |
| FbsH WT | 3-Amino-Sal | 1 | $1.35 \pm 0.05$ | $125 \pm 15.1$ | $(2.25 \pm 0.08) \times 10^{-2}$ | $1.81 \times 10^2$ |
| FbsH WT | 3-Methyl-Sal | 1 | $2.21 \pm 0.04$ | $72.7 \pm 5.06$ | $(3.68 \pm 0.07) \times 10^{-2}$ | $5.06 \times 10^2$ |
| FbsH WT | 4-Fluoro-Sal | 1 | $0.71 \pm 0.04$ | $133 \pm 24$ | $(1.18 \pm 0.07) \times 10^{-2}$ | $8.94 \times 10^1$ |
| FbsH WT | 4-Amino-Sal | 5 | $0.69 \pm 0.07$ | $630 \pm 122$ | $(0.23 \pm 0.02) \times 10^{-2}$ | $3.65 \times 10^0$ |
| FbsH WT | 4-Methyl-Sal | 5 | $1.01 \pm 0.07$ | $366 \pm 75$ | $(0.34 \pm 0.02) \times 10^{-2}$ | $9.16 \times 10^0$ |
| FbsH WT | 4-Azido-Sal | 5 | --<sup>a</sup> | -- | -- | -- |
| FbsH WT | 4-Nitro-Sal | 5 | --<sup>a</sup> | -- | -- | -- |
| FbsH F350A | Sal | 1 | $0.58 \pm 0.02$ | $22.7 \pm 3.80$ | $(9.70 \pm 0.40) \times 10^{-3}$ | $4.28 \times 10^2$ |
| FbsH F350A | 3-Bromo-Sal | 0.8 | $0.50 \pm 0.04$ | $51.6 \pm 10.2$ | $(1.04 \pm 0.07) \times 10^{-2}$ | $2.01 \times 10^2$ |
| FbsH F350A | 4-Fluoro-Sal | 1 | $0.35 \pm 0.02$ | $17.7 \pm 4.33$ | $(5.83 \pm 0.36) \times 10^{-3}$ | $3.26 \times 10^2$ |
| FbsH F350A | 4-Amino-Sal | 2 | $0.59 \pm 0.03$ | $25.8 \pm 5.51$ | $(4.93 \pm 0.27) \times 10^{-3}$ | $1.92 \times 10^2$ |
| FbsH F350A | 4-Methyl-Sal | 2 | $0.76 \pm 0.04$ | $95.5 \pm 20.1$ | $(6.32 \pm 0.35) \times 10^{-3}$ | $6.62 \times 10^1$ |

<sup>a</sup>No detectable activity

### Mutagenesis of Active Site to Expand Substrate Promiscuity

Examining the binding pocket residues of FbsH, the C4 carbon of the DHB or salicylic acid group is positioned 3.5 Å from the Cγ of Val340, 3.6 Å from the Cβ of Ser251, and 4.2 Å from the Cγ of Val348, suggesting that steric interactions may limit the ability of the C4-substituted substrates from adopting a proper position for efficient adenylation. Considering the entire pocket, within 5 Å of the ligand are His244, Asn246, Phe247, Ser250, Ser251, Val340, and Val348 (Figure 2C). His244 is part of the conserved A4 motif^22^ that positions the carboxylate for the adenylation reaction. Phe247, which is replaced with a Tyr in the other structurally characterized aryl adenylating domains, appears to form a stacking arrangement with the substrate. Additionally, Asn246 and Ser251 interact with hydroxyl groups on the DHB substrate and were therefore left in place.

We therefore focused our attention on Val340 and Val348, which form a hydrophobic base of the pocket. To expand the pocket to accommodate larger substituents added onto C4, such as 4-azido salicylic acid, we also targeted Phe350, which lies below Val340 and Val348 and is 6Å from the C4 position of the DHB ligand and could therefore be considered a second shell residue. Removal of the Phe350 side chain could directly expand the active site or may enable movement of residues that are positioned closer to the aryl substrate. Val340, Val348, and Phe350 were all replaced with alanine. The V340A mutant protein failed to express in soluble form instead resulting in the formation of insoluble inclusion bodies under a variety of heterologous expression conditions.

The V348A and F350A mutants were next screened for specific activity with a panel of substrates (Figure 3A) showing that the F350A mutant improved activity with the 4-substituted analogs, as well as a few 3-substituted analogs. In contrast, the V348A mutant showed diminished adenylating activity towards the analogs, suggesting that Val348 may form hydrophobic interactions with the substrates that are important for binding and/or orientating the substrate for acyl adenylate formation.

We collected steady-state kinetic data to better understand the enhancement in activity that was observed with the FbsH F350A mutant with the DHB analogs (Table 1, Supplemental Figure 4). Salicylic acid and 3-bromosalicylic acid produced kinetic parameters that were comparable for the wild-type and mutant enzymes, with minimal changes in apparent k_cat_/K_M_. However, significant changes in catalytic efficiency of the FbsH F350A mutant relative to WT FbsH were observed for the C4-substituted analogs that were tested. For 4-fluorosalicylic acid, an ∼8-fold decrease in apparent K_M_ was observed resulting in a ∼4-fold increase in catalytic efficiency. With 4-methylsalicylic acid, a similar ∼4-fold reduction in apparent K_M_ and a ∼2-fold improvement in apparent k_cat_ resulted in the apparent k_cat_/K_M_ improving ∼7-fold. The most dramatic improvement in catalytic efficiency was observed with the substrate 4-amino salicylic acid, where a ∼25-fold decline in apparent K_M_ combined with a ∼2-fold improvement in apparent k_cat_ resulted in a ∼52-fold increase in catalytic efficiency. This improved K_M_ of 26 µM is even comparable to the wild-type FbsH K_M_ of 21 µM with the native substrate DHB. The observed significant improvements in catalytic efficiencies indicate that Phe350 in FbsH influences the binding affinity and adenylation rate of 4-substituted benzoate substrate analogs.

Overall, our steady-state kinetic data indicate that an expanded binding pocket of the FbsH aryl acid adenylation domain can enhance the promiscuity for binding and adenylating substrate analogs. Here, we specifically targeted the base of the pocket, near the C4 position of the aryl ring. Interestingly, despite being a relatively minor change, removal of the two methyl groups of Val340 appeared to influence protein solubility. Further, mutation of Val348 to alanine resulted in a reduction in specific activity in all substrates. We propose that this residue, which forms one hydrophobic wall of the binding pocket, may support positioning of the aromatic ring for adenylation. Most interesting, mutation of the second shell residue Phe350 improves the apparent catalytic efficiency of the enzyme to allow activity with several C4 substituted analogs, presumably by providing room at the base of the pocket.

### Full *in vitro* Reconstitution Assay to Monitor Intermediates and Product Formation

We have previously reconstituted fimsbactin biosynthesis using the core NRPS enzymes, FbsEFGH, and the building blocks DHB, serine, and ahPut.^44^ We therefore examined the ability to produce fimsbactin analogs from alternate aromatic building blocks (Figure 4A). Additionally, as the adenylation domains of NRPS modules are the gatekeepers for substrate activation and tethering to peptidyl carrier proteins via the phosphopantetheinyl thioesters, we also asked whether the mutant FbsH enzymes might enhance the diversity of fimsbactin variants that could be produced.

**Figure 4:**
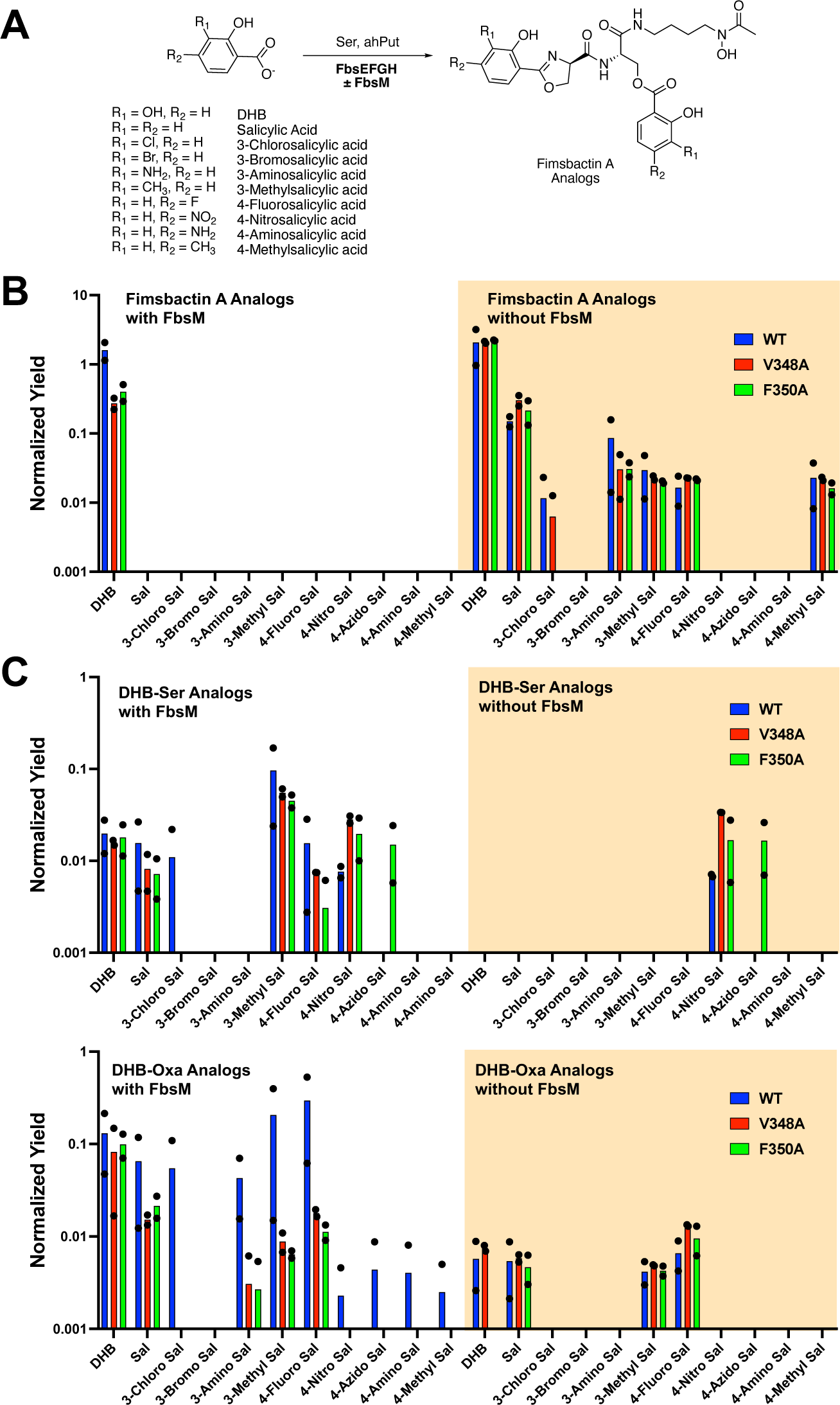
Full *in vitro* reconstitution of Fimsbactin with wild-type and mutant FbsH and DHB analogs. (A) Structures of Fimsbactin A analogs produced from numerous DHB analogs. (B) Normalized yields of Fimsbactin and Fimsbactin analogs, with and without FbsM. Data is plotted as an average of two replicates performed on separate days. Black dots represent individual replicates. (C) Normalized yields of DHB-Ser (top) and DHB-oxa (bottom) analogs, with and without FbsM. Data is plotted as an average of two replicates performed on separate days. Black dots represent individual replicates. Details of normalization and peak validation are provided in the methods section.

We explored combinations of FbsH WT, V348A mutant, or F350A mutant with FbsE, FbsF, and FbsG, in the presence or absence of the editing thioesterase FbsM under *in vitro* reaction conditions using LCMS to detect biosynthetic intermediates and product analogs. Reactions were performed as biological replicates in two independent trials. We monitored the formation of analogs of two intermediates, aryl-serine and aryl-oxazoline, as well as analogs of fimsbactin A that were produced in the reconstitution reaction (Supplemental Figures 5 and 6). Considering the ability of thioesterase FbsM to act as a proofreading enzyme that can clear misloaded amino acids and hydrolyze biosynthetic intermediates from numerous PCP domains,^44^ we carried out the assay both in the presence and absence of FbsM to test if FbsM changes the product pools.

Although FbsH shows similar apparent kinetics with DHB and salicylic acid, the full reconstitution reaction shows that, in the absence of FbsM, production of the salicyl analog of fimsbactin A lacking the 3-OH group on both aryl rings is reduced by an order of magnitude (Figure 4B). This suggests that a downstream step in the reaction, after loading of the aryl-carrier protein domain of FbsE, serves as a checkpoint to limit throughput to the fimsbactin analog.

We then examined the panel of salicylic acid analogs used in the FbsH adenylation reactions. Relative to salicylic acid, the full throughput to fimsbactin analogs is reduced by an additional 5-10 fold with five of the nine analogs tested (Figure 4B), with no observable peak for 3-bromosalicylate and three of the C4-substituted salicylate analogs. This suggests that the downstream catalytic domains also prefer the presence of both the hydroxyl group on the aryl ring and the lack of a C4 modification. In the presence of FbsM, the only fimsbactin analog produced is the native product formed from 2,3-DHB; fimsbactin analogs built from all remaining substrates could not be detected. As the FbsH adenylating activity is comparable for the many analogs, this suggests that the loaded aryl-carrier protein domain, or a subsequent downstream intermediate, reacts slowly enough that the FbsM enzyme can cleave the loaded thioester releasing the peptide intermediate in competition with the full pathway.

To examine this, we monitored the formation of the DHB-Ser and DHB-Oxa intermediates in the presence and absence of FbsM. We note that the DHB-Ser *m/z* value is identical for both *N*-DHB-Ser and *O-*DHB-Ser aryl amino acids and no effort was made to distinguish the two compounds. Both in the presence and absence of FbsM, a significant amount of the aryl-Ser and aryl-Oxa intermediates is formed (Figure 4C), suggesting that the slow reaction progress makes the intermediates available for catalytic “proof-reading” release by FbsM. The presence of FbsM did increase the variety of aryl-Ser (7/11 substrates) and aryl-Oxa (10/11) intermediates compared to reactions lacking FbsM (2/11 for aryl-Ser; 4/11 for aryl-Oxa). Interestingly, even with the true substrate DHB, significant amounts of the DHB-Ser and DHB-Oxa intermediates are released, suggesting that the thioesterase FbsM still catalyzes the release of limited amounts of the peptide intermediates even going at full speed with the native substrate,

## Discussion

Here, we expand on prior studies to explore NRPS adenylation domain promiscuity.^35, 36, 47^ In particular, we have focused on a complex, branching NRPS siderophore assembly that incorporates two aryl caps at both the N-terminus of the peptide as well as at a Ser hydroxyl group as a branching ester via acylation (Figure 1). This study expands our understanding of determinants of NRPS substrate promiscuity and further interrogates the ability of alternate starter units to progress through the entire NRPS pathway. Our studies suggest downstream checkpoint domains within the pathway that will inform subsequent studies to increase the promiscuity of additional catalytic domains.

Using X-ray crystallography, we solved new structures of the standalone adenylation domain FbsH bound to unique ligands. As this “gatekeeper” domain acts as a point of entry for aryl acids like DHB, we explored its promiscuity by feeding it different substrate analogs. Using structural analysis, we engineered mutants to enhance the ability of FbsH to recognize non-native substrates and identified determinants of its substrate selectivity for alternate substrates. We propose that removal of the aromatic ring of Phe350, located at the base of its substrate binding pocket, enables the enzyme to accommodate 4-substituted substrate analogs of DHB, with limited impact on the 3-substituted analogs that we tested. Additionally, the Val348 residue appears to contribute important hydrophobic interactions to properly orient the analogs, and any increase in the size in the cavity is offset by this effect. Our preliminary substrate screening data supports this idea as we see an improvement in activity only for analogs with the F350A mutant whereas a decline in activity for the analogs with the V348A mutant. Moreover, our steady-state kinetic data further support our proposal on these specificity determinants, particularly Phe350. With wild-type FbsH, we observe catalytic efficiencies to be typically higher for 3-substituted analogs compared to 4-substituted analogs. When compared to the F350A mutant of FbsH, the catalytic efficiencies are significantly higher for these 4-substituted analogs, with a stark 52-fold improvement for 4-amino salicylic acid. We propose that the larger steric size of the Phe350 side chain likely causes a steric block with the 4^th^ position substituents (Figure 2b), which is not present when Phe350 is substituted with a significantly smaller alanine, resulting in this dramatic difference in activity. Furthermore, our findings with the FbsH mutants are consistent with previous studies where NRPS adenylation domains were engineered to have an expanded binding pocket to tolerate substrate analogs of DHB.^35, 36^

Previous efforts with EntE had primarily focused on the Asn235 residue, which is responsible for the recognition of DHB as it forms hydrogen bonds with the hydroxyl groups of DHB. Once this asparagine was mutated to glycine, the EntE mutant with an expanded binding pocket had improved activity with several C2- and C3-substituted benzoic acid analogs.^35^ Furthermore, EntE has also been engineered to swap its NRPS code from a primarily DHB-binding to a Sal-binding adenylation domain, targeting first shell residues equivalent to tyr247, Asn246, Ser251, Val348, and Asn349, resulting in these mutations also having a higher activity with 2 and 3-substituted benzoic acid analogs when compared to wild-type.^36^ In contrast, as our focus was on expanding activity for C3 and C4-substituted salicylic acid analogs; therefore, we primarily focused on the base of the pocket, while retaining the N246 residue to allow proper binding and recognition of the hydroxyl group of the salicylic acid analogs.^36^ These EntE mutants also had higher activity with 2 and 3-substituted benzoic acid analogs when compared to wild-type. Combining the results of this study and previous mutations in EntE, future studies could utilize the EntE mutations to switch the FbsH NRPS code to a Sal binding enzyme in combination with mutants that expand the pocket at its base to generate an improved enzyme with comparable catalytic efficiencies for C4-substituted salicylic analogs compared to DHB for wild-type FbsH.

Our expanded understanding of FbsH substrate selectivity encouraged us to further explore downstream NRPS selectivity in the fimsbactin biosynthetic pathway. Previously, we proposed a mechanism of fimsbactin assembly by several multidomain NRPS enzymes in a unique non-linear fashion, establishing a foundation for understanding downstream selectivity for diverse substrate analogs.^44^ Data obtained from our full reconstitution experiments showed downstream selectivity in the pathway, as well as the role of the type 2 thioesterase FbsM in proofreading against non-cognate substrates. However, the production of diverse intermediate products from these analogs demonstrates that these aryl acid derivatives can be carried through the entire NRPS assembly line. We observed the presence of several DHB-Ser and DHB- Oxa analogs, indicating that the aryl acid analogs were carried through the primary adenylation reaction by the adenylation domain of FbsH, followed by the condensation reaction to the serine residue. Some analogs were further cyclized by the cyclization domain of FbsE to produce DHB-Oxa analogs. Notably, in the absence of FbsM, we observe the presence of several fimsbactin A analogs suggesting that some DHB analogs were processed through the entire NRPS assembly line where FbsG catalyzed the final aminolysis step using *N-*acetyl-*N*-hydroxyputrescine (ahPut) as the cleaving nucleophile to form the complete fimsbactin A analogs. The accumulation of intermediates in the presence of FbsM shows the additional domain engineering is required to further improve the yield of the fimsbactin A analogs. Recent studies from other groups have shown that condensation domains represent a common bottleneck in NRPS assembly lines.^48-52^

The production of siderophore analogs could lead to novel siderophore-based antimicrobials that possess activity against antibiotic-resistant bacteria such as *A. baumannii*. Our work lays a foundation for the generation of fimsbactin derivatives with antimicrobial activity that could be used as building blocks for conjugation with existing antibiotics. In addition, our work supports chemoenzymatic approaches for siderophore-based antibiotic production, where enzymes substitute complex chemical organic synthesis, making drug discovery cheaper, more efficient, and more environmentally friendly. Based on our discoveries, we propose that engineering enzymes will be a crucial first step for taking chemoenzymatic approaches to create a more effective drug discovery pipeline.

## Materials and Methods

### Cloning and gene annotation

Cloned wild-type *fbsH* in the pET28a vector containing the His-tag was obtained for heterologous expression in *E. coli* (GenBank Assembly Accession: WP_049594809.1).^44^ FbsH variants were generated using site-directed mutagenesis using PCR. Oligonucleotides used for mutagenesis are shown in Supplemental Table 3. The codon-optimized gene and FbsH protein sequence are presented in Supplemental Tables 4 and 5.

### Recombinant protein expression and purification

Wild-type FbsH in the pET-28a vector was transformed into *E. coli* BL21 DE3 cells. Cells were grown overnight in 50 mL LB containing 50 ug/ml kanamycin at 37°C. 10 mL of overnight culture was then inoculated into 1 L LB medium supplemented with 50 µg/ml kanamycin. Cultures were grown at 37°C with agitation until the OD_600_ reached 0.6-0.8 when they were cooled at 4°C for 10 min. Protein expression was induced with 0.5 mM IPTG and cultures were grown for 18 hours at 20°C. Cells were cooled at 4°C and harvested via centrifugation at 6000 rpm for 10 min and frozen at -80°C. All subsequent purification steps were carried out at 4°C.

His-tagged FbsH was purified using immobilized metal affinity chromatography. Cells were resuspended in lysis buffer (50 mM HEPES pH 7.5, 500 mM NaCl, 0.2 mM TCEP, 20 mM imidazole, and 10% glycerol) and lysed using sonication (cycles of 5 sec on, 10 sec off for a total on time of 5 min at 60% amplitude). The resulting cell lysate was centrifuged at 40,000 rpm for 40 min. The supernatant was decanted and filtered using a 0.45 µm filter. The filtrate was passed over a 5 ml nickel-loaded HisTrap column. The column was washed with 10 column volumes of lysis buffer followed by 5 column volumes of 6% elution buffer (50 mM HEPES pH 7.5, 500 mM NaCl, 0.2 mM TCEP, 300 mM imidazole, and 10% glycerol). The immobilized protein was then eluted with 15 column volumes of elution buffer (gradient) and dialyzed overnight with dialysis buffer (20 mM HEPES pH 7.5, 100 mM NaCl, and 0.2 mM TCEP). The dialyzed protein was collected the next day and concentrated using a 15 mL Amicon Centrifugation filter (30 kDa MWCO) until the concentration reached 10-15 mg/ml. The concentrated protein was centrifuged at 14000 rpm to remove aggregates. Protein to be used in kinetic studies was flash-frozen in liquid nitrogen and stored at -80°C. For structural studies, the protein was further purified using size exclusion chromatography for crystallization trials. The concentrated protein was run over gel filtration buffer (20 mM HEPES pH 7.5, 100 mM NaCl, and 0.2 mM TCEP) on a HiPrep 26/60 Sephacryl S-300 column. The purified protein was concentrated again until it reached 10-20 mg/ml and centrifuged at 14000 rpm to remove any aggregates. Purified tagged FbsH was then flash-frozen in liquid nitrogen and stored at -80°C. The same protocol was used for the FbsH mutants.

### Crystallization and structure solution

His-tagged FbsH purified via nickel affinity and size exclusion chromatography was used for crystallographic trials. A sparse matrix crystal screen was performed with a Gryphon robot at 14°C with 13 mg/ml FbsH in the presence of 600 µM salicyl-AMS inhibitor. Initial crystal hits of 13 mg/ml FbsH were observed within two days of setting up trays, in the presence of 20% PEG 3350 and several salts. After optimization, crystals were obtained at 0.1 M HEPES pH 7.5 and 12 – 15% PEG 3350. The final crystals used for diffraction experiments were obtained from 12 % PEG 3350, 0.1 M HEPES, pH 7.5, and 0.5 mM DHB and AMP for the DHB-bound structure, or with 12 % PEG 3350, 0.1 M HEPES, pH 7.5, 600 µM Sal-AMS for the inhibitor structure. Crystals were harvested within a week of observation cryo-protected with 16% ethylene glycol in crystallization cocktail, and flash-frozen in liquid nitrogen.

Diffraction data for FbsH single crystals was remotely collected at APS using the beamline 23IDD. Both datasets (one bound to DHB or Sal-AMS) were processed at 2.2Å using Xia2/Dials in the *P*1 space group. The Sal-AMS structure was solved via molecular replacement with Phaser in the Phenix Suite using an AlphaFold model^46^ of FbsH as the search model. The Sal-AMS model lacking ligands was in turn used for solution of the DHB structure. Because of the conformational dynamics of the C-terminal subdomain, the search model consisted of only the N-terminal subdomain (residues 1-439). This resulted in solutions for each dataset containing two molecules per asymmetric unit. The C-terminal subdomain for each monomer had weaker densities and was manually built. Manual model building and refinement were carried out using Phenix.refine^53^ and COOT.^54^

### Analysis of FbsH adenylation activity via the NADH Consumption Assay

A hydroxamate-coupled NADH assay was utilized to measure the apparent kinetic constants derived from activity of FbsH wild-type and mutants.^55, 56^ The reaction mixture contained 100 mM HEPES pH 7.5, 15 mM MgCl_2_, 3 mM ATP, 50 mM hydroxylamine, 3 mM phosphoenolpyruvate, 0.8 mM NADH, 10 Units/ml of each coupling enzyme myokinase, pyruvate kinase, lactate dehydrogenase, and varying concentrations of substrates and FbsH wild-type and mutant proteins. For initial measurement of specific activity to screen substrates, 500 µM of each substrate and 5 µM enzyme was used whereas for steady-state kinetic analysis, this varied depending on the substrate and enzyme being tested (typically 0.5-5 µM enzyme and 6-8 different substrate concentrations per assay). The assay was conducted in 100 µL reaction volume in clear 96-well polystyrene plates. All reactions were done in triplicate. The master mix was first incubated for 10 min at 37°C without FbsH or substrate. In a separate plate, 10 µL of protein and substrate were added without contacting each other. Once the master mix was incubated, 80 µL of the master mix was added to wells containing protein and substrate. Absorbance at 340 nm was continuously monitored for 20 minutes at 37°C using a Biotek Citation 1 plate reader. Absorbance at 340 nm was later converted to concentrations of NADH using the molar extinction coefficient of NADH (6220 M^-1^cm^-1^). The rate of decay of NADH was used to generate specific activity plots for substrate screening and Michaelis-Menten plots to generate steady-state kinetic constants using GraphPad Prism 10.

### Phosphopantetheinylation of FbsE, FbsF, and FbsG

In a 1.5 mL Eppendorf tube, 25 μM of *apo*-FbsE, *apo*-FbsF, or *apo*-FbsG (wild-type or mutants) was incubated in 50 mM HEPES buffer at pH 7.5 with 10 mM MgCl_2_, 100 μM CoASH. and 1 μM the promiscuous phosphopantetheinyl transferase, Sfp. Reactions were left at room temperature for 2 h, and then aliquoted and stored at -80 °C for future use as the source of *holo*-FbsE, *holo*-FbsF, and *holo*-FbsG.

### Chemoenzymatic synthesis of fimsbactin A, analogs, and intermediates

Reactions were performed at 100 μL total working volume by adapting the previously reported protocol.^44^ Full reactions contained 5 mM ATP, 1 mM aryl acid, 1 mM L-Ser, 0.5 mM ahPut, 1 μM *holo*-FbsE, 1 μM *holo*-FbsF, 1 μM *holo*-FbsG, 1 μM FbsH (wild type or mutant), and 50 mM HEPES at pH 7.5. Reactions were performed at room temperature. At 3 h time points, the reactions were quenched by the addition of 400 μL of MeOH containing 120 μM Fmoc-Ala as an internal standard. Mixtures were centrifuged for 2 min at 13,000 r.p.m. to remove precipitated enzyme, and the supernatants were analyzed by LC-MS using a gradient of 5% B to 95% B over 20 min. Ion counts corresponding to the expected *m/z* value of the [M+H]^+^ molecular ion of fimsbactin A, E, F, DHB-Ser, and DHB-Oxa, along with analogs derived from various aryl acid substrates, were extracted from total ion chromatograms acquired in positive ion mode. Reactions were run in two biological replicates, performed on different days, and compared to a standard reaction in which no enzymes were added. To identify the peaks (Supplemental Figures 5 and 6), we developed a set of criteria that were used for validation. First, the molecular ion must be observed in the reaction trace and not observed in the no enzyme control. Second, the retention time is unique to the observed molecular ion. Third, the retention time is reasonable based on fimsbactin standard compounds and intermediates with DHB. Finally, the retention time does not overlap directly with another compound that can undergo in-source fragmentation in the mass spectrometer to generate the molecular ion of interest. The extracted ion counts for each analyte were normalized to the Fmoc-Ala internal standard by dividing the peak height of the analyte’s extracted ion to the peak height corresponding to the extracted ion for Fmoc-Ala.

## Supporting Information

Supporting Information is available for this article.

## Authorship Contributions

The experimental work described in the manuscript was performed by S.F.A., A.B., and J.Y. under the supervision of T.A.W. and A.M.G. The manuscript was written by S.F.A., T.A.W., and A.M.G. All authors edited and reviewed the final manuscript.

## Data Availability

The structures of FbsH have been deposited with the worldwide Protein Data Bank (FbsH bound to 2,3-dihydroxybenzoic acid, **9BHY**; FbsH bound to Sal-AMS, PDB **9BHZ**)

## Acknowledgments

The investigation was supported by NIH grants GM136235 (to AMG), NSF CAREER 1654611 (to TAW), and the Washington University College of Arts & Sciences Incubator for Transdisciplinary Futures. Diffraction data were collected at APS (Advanced Photon Source, Lemont, IL 60439, USA). GM/CA@APS has been funded by the National Cancer Institute (ACB-12002) and the National Institute of General Medical Sciences (AGM-12006, P30GM138396). This research used resources from the Advanced Photon Source, a U.S. Department of Energy (DOE) Office of Science User Facility operated for the DOE Office of Science by Argonne National Laboratory under Contract No. DE-AC02-06CH11357. AB is supported by an NSF GRFP fellowship (DGE 2139839). We thank Professor Courtney Aldrich (University of Minnesota) for the kind gift of the Sal-AMS inhibitor.

## References

(1) Hider, R. C.; Kong, X. Chemistry and biology of siderophores. Nat Prod Rep 2010, 27 (5), 637–657.

(2) Kadner, R. J.; McElhaney, G. Outer membrane-dependent transport systems in Escherichia coli: turnover of TonB function. J Bacteriol 1978, 134 (3), 1020–1029.

(3) Faraldo-Gomez, J. D.; Sansom, M. S. Acquisition of siderophores in gram-negative bacteria. Nat Rev Mol Cell Biol 2003, 4 (2), 105–116.

(4) Cook-Libin, S.; Sykes, E. M. E.; Kornelsen, V.; Kumar, A. Iron Acquisition Mechanisms and Their Role in the Virulence of Acinetobacter baumannii. Infect Immun 2022, 90 (10), e0022322.

(5) Artuso, I.; Poddar, H.; Evans, B. A.; Visca, P. Genomics of Acinetobacter baumannii iron uptake. Microb Genom 2023, 9 (8).

(6) Lee, Y. R.; Yeo, S. Cefiderocol, a New Siderophore Cephalosporin for the Treatment of Complicated Urinary Tract Infections Caused by Multidrug-Resistant Pathogens: Preclinical and Clinical Pharmacokinetics, Pharmacodynamics, Efficacy and Safety. Clin Drug Investig 2020, 40 (10), 901–913.

(7) Runci, F.; Gentile, V.; Frangipani, E.; Rampioni, G.; Leoni, L.; Lucidi, M.; Visaggio, D.; Harris, G.; Chen, W.; Stahl, J.;, et al. Contribution of Active Iron Uptake to Acinetobacter baumannii Pathogenicity. Infect Immun 2019, 87 (4).

(8) Hamidian, M.; Hall, R. M. Dissemination of novel Tn7 family transposons carrying genes for synthesis and uptake of fimsbactin siderophores among Acinetobacter baumannii isolates. Microb Genom 2021, 7 (3).

(9) Klebba, P. E.; Newton, S. M. C.; Six, D. A.; Kumar, A.; Yang, T.; Nairn, B. L.; Munger, C.; Chakravorty, S. Iron Acquisition Systems of Gram-negative Bacterial Pathogens Define TonB-Dependent Pathways to Novel Antibiotics. Chem Rev 2021, 121 (9), 5193–5239.

(10) Lamb, A. L. Breaking a pathogen’s iron will: Inhibiting siderophore production as an antimicrobial strategy. Biochim Biophys Acta 2015, 1854 (8), 1054–1070.

(11) Carroll, C. S.; Moore, M. M. Ironing out siderophore biosynthesis: a review of non-ribosomal peptide synthetase (NRPS)-independent siderophore synthetases. Crit Rev Biochem Mol Biol 2018, 53 (4), 356–381.

(12) Staunton, J.; Wilkinson, B. Combinatorial biosynthesis of polyketides and nonribosomal peptides. Curr Opin Chem Biol 2001, 5 (2), 159–164.

(13) Miller, B. R.; Gulick, A. M. Structural Biology of Nonribosomal Peptide Synthetases. Methods Mol Biol 2016, 1401, 3–29.

(14) Patel, K. D.; MacDonald, M. R.; Ahmed, S. F.; Singh, J.; Gulick, A. M. Structural advances toward understanding the catalytic activity and conformational dynamics of modular nonribosomal peptide synthetases. Nat Prod Rep 2023, 40 (9), 1550–1582.

(15) Patel, K. D.; Ahmed, S. F.; MacDonald, M. R.; Gulick, A. M. Structural Studies of Modular Nonribosomal Peptide Synthetases. Methods Mol Biol 2023, 2670, 17–46.

(16) Miyanaga, A.; Kudo, F.; Eguchi, T. Recent advances in the structural analysis of adenylation domains in natural product biosynthesis. Curr Opin Chem Biol 2022, 71, 102212.

(17) Dekimpe, S.; Masschelein, J. Beyond peptide bond formation: the versatile role of condensation domains in natural product biosynthesis. Nat Prod Rep 2021, 38 (10), 1910–1937.

(18) Horsman, M. E.; Hari, T. P.; Boddy, C. N. Polyketide synthase and non-ribosomal peptide synthetase thioesterase selectivity: logic gate or a victim of fate? Nat Prod Rep 2016, 33 (2), 183–202.

(19) Patel, H. M.; Tao, J.; Walsh, C. T. Epimerization of an L-cysteinyl to a D-cysteinyl residue during thiazoline ring formation in siderophore chain elongation by pyochelin synthetase from Pseudomonas aeruginosa. Biochemistry 2003, 42 (35), 10514–10527.

(20) Ronnebaum, T. A.; McFarlane, J. S.; Prisinzano, T. E.; Booker, S. J.; Lamb, A. L. Stuffed Methyltransferase Catalyzes the Penultimate Step of Pyochelin Biosynthesis. Biochemistry 2019, 58 (6), 665–678.

(21) Wuest, W. M.; Sattely, E. S.; Walsh, C. T. Three siderophores from one bacterial enzymatic assembly line. J Am Chem Soc 2009, 131 (14), 5056–5057.

(22) Gulick, A. M. Conformational dynamics in the Acyl-CoA synthetases, adenylation domains of non-ribosomal peptide synthetases, and firefly luciferase. ACS Chem Biol 2009, 4 (10), 811–827.

(23) Stachelhaus, T.; Mootz, H. D.; Marahiel, M. A. The specificity-conferring code of adenylation domains in nonribosomal peptide synthetases. Chem Biol 1999, 6 (8), 493–505.

(24) Challis, G. L.; Ravel, J.; Townsend, C. A. Predictive, structure-based model of amino acid recognition by nonribosomal peptide synthetase adenylation domains. Chem Biol 2000, 7 (3), 211–224.

(25) Rusnak, F.; Faraci, W. S.; Walsh, C. T. Subcloning, expression, and purification of the enterobactin biosynthetic enzyme 2,3-dihydroxybenzoate-AMP ligase: demonstration of enzyme-bound (2,3-dihydroxybenzoyl)adenylate product. Biochemistry 1989, 28 (17), 6827–6835.

(26) Heard, S. C.; Winter, J. M. Structural, biochemical and bioinformatic analyses of nonribosomal peptide synthetase adenylation domains. Nat Prod Rep 2024, 41 (7), 1180–1205.

(27) May, J. J.; Kessler, N.; Marahiel, M. A.; Stubbs, M. T. Crystal structure of DhbE, an archetype for aryl acid activating domains of modular nonribosomal peptide synthetases. Proc Natl Acad Sci U S A 2002, 99 (19), 12120–12125.

(28) Shelton, C. L.; Meneely, K. M.; Ronnebaum, T. A.; Chilton, A. S.; Riley, A. P.; Prisinzano, T. E.; Lamb, A. L. Rational inhibitor design for Pseudomonas aeruginosa salicylate adenylation enzyme PchD. J Biol Inorg Chem 2022, 27 (6), 541–551.

(29) Sundlov, J. A.; Gulick, A. M. Structure determination of the functional domain interaction of a chimeric nonribosomal peptide synthetase from a challenging crystal with noncrystallographic translational symmetry. Acta Crystallogr D Biol Crystallogr 2013, 69 (Pt 8), 1482–1492.

(30) Sundlov, J. A.; Shi, C.; Wilson, D. J.; Aldrich, C. C.; Gulick, A. M. Structural and functional investigation of the intermolecular interaction between NRPS adenylation and carrier protein domains. Chem Biol 2012, 19 (2), 188–198.

(31) Tan, X. F.; Dai, Y. N.; Zhou, K.; Jiang, Y. L.; Ren, Y. M.; Chen, Y.; Zhou, C. Z. Structure of the adenylation-peptidyl carrier protein didomain of the Microcystis aeruginosa microcystin synthetase McyG. Acta Crystallogr D Biol Crystallogr 2015, 71 (Pt 4), 873–881.

(32) Alexander, E. M.; Kreitler, D. F.; Guidolin, V.; Hurben, A. K.; Drake, E.; Villalta, P. W.; Balbo, S.; Gulick, A. M.; Aldrich, C. C. Biosynthesis, Mechanism of Action, and Inhibition of the Enterotoxin Tilimycin Produced by the Opportunistic Pathogen Klebsiella oxytoca. ACS Infect Dis 2020, 6 (7), 1976–1997.

(33) Vergnolle, O.; Xu, H.; Tufariello, J. M.; Favrot, L.; Malek, A. A.; Jacobs, W. R., Jr.; Blanchard, J. S. Post-translational Acetylation of MbtA Modulates Mycobacterial Siderophore Biosynthesis. J Biol Chem 2016, 291 (42), 22315–22326.

(34) Drake, E. J.; Duckworth, B. P.; Neres, J.; Aldrich, C. C.; Gulick, A. M. Biochemical and structural characterization of bisubstrate inhibitors of BasE, the self-standing nonribosomal peptide synthetase adenylate-forming enzyme of acinetobactin synthesis. Biochemistry 2010, 49 (43), 9292–9305.

(35) Ishikawa, F.; Miyanaga, A.; Kitayama, H.; Nakamura, S.; Nakanishi, I.; Kudo, F.; Eguchi, T.; Tanabe, G. An Engineered Aryl Acid Adenylation Domain with an Enlarged Substrate Binding Pocket. Angew Chem Int Ed Engl 2019, 58 (21), 6906–6910.

(36) Ishikawa, F.; Nohara, M.; Nakamura, S.; Nakanishi, I.; Tanabe, G. Precise Probing of Residue Roles by NRPS Code Swapping: Mutation, Enzymatic Characterization, Modeling, and Substrate Promiscuity of Aryl Acid Adenylation Domains. Biochemistry 2020, 59 (4), 351–363.

(37) Ishikawa, F.; Kitayama, H.; Nakamura, S.; Takashima, K.; Nakanishi, I.; Tanabe, G. Activity, Binding, and Modeling Studies of a Reprogrammed Aryl Acid Adenylation Domain with an Enlarged Substrate Binding Pocket. Chem Pharm Bull (Tokyo) 2021, 69 (2), 222–225.

(38) Tripathi, A.; Park, S. R.; Sikkema, A. P.; Cho, H. J.; Wu, J.; Lee, B.; Xi, C.; Smith, J. L.; Sherman, D. H. A Defined and Flexible Pocket Explains Aryl Substrate Promiscuity of the Cahuitamycin Starter Unit-Activating Enzyme CahJ. Chembiochem 2018, 19 (15), 1595–1600.

(39) Zhang, K.; Kries, H. Biomimetic engineering of nonribosomal peptide synthesis. Biochem Soc Trans 2023, 51 (4), 1521–1532.

(40) Katsuyama, Y.; Miyanaga, A. Recent advances in the structural biology of modular polyketide synthases and nonribosomal peptide synthetases. Curr Opin Chem Biol 2022, 71, 102223.

(41) Proschak, A.; Lubuta, P.; Grun, P.; Lohr, F.; Wilharm, G.; De Berardinis, V.; Bode, H. B. Structure and biosynthesis of fimsbactins A-F, siderophores from Acinetobacter baumannii and Acinetobacter baylyi. Chembiochem 2013, 14 (5), 633–638.

(42) Bohac, T. J.; Fang, L.; Banas, V. S.; Giblin, D. E.; Wencewicz, T. A. Synthetic Mimics of Native Siderophores Disrupt Iron Trafficking in Acinetobacter baumannii. ACS Infect Dis 2021, 7 (8), 2138–2151.

(43) Bohac, T. J.; Fang, L.; Giblin, D. E.; Wencewicz, T. A. Fimsbactin and Acinetobactin Compete for the Periplasmic Siderophore Binding Protein BauB in Pathogenic Acinetobacter baumannii. ACS Chem Biol 2019, 14 (4), 674–687.

(44) Yang, J.; Wencewicz, T. A. In Vitro Reconstitution of Fimsbactin Biosynthesis from Acinetobacter baumannii. ACS Chem Biol 2022, 17 (10), 2923–2935.

(45) Qiao, C.; Gupte, A.; Boshoff, H. I.; Wilson, D. J.; Bennett, E. M.; Somu, R. V.; Barry, C. E., 3rd; Aldrich, C. C. 5’-O-[(N-acyl)sulfamoyl]adenosines as antitubercular agents that inhibit MbtA: an adenylation enzyme required for siderophore biosynthesis of the mycobactins. J Med Chem 2007, *50* (24), 6080-6094.

(46) Jumper, J.; Evans, R.; Pritzel, A.; Green, T.; Figurnov, M.; Ronneberger, O.; Tunyasuvunakool, K.; Bates, R.; Zidek, A.; Potapenko, A.;, et al. Highly accurate protein structure prediction with AlphaFold. Nature 2021, 596 (7873), 583–589.

(47) Hegde, P.; Orimoloye, M. O.; Sharma, S.; Engelhart, C. A.; Schnappinger, D.; Aldrich, C. C. Polyfluorinated salicylic acid analogs do not interfere with siderophore biosynthesis. Tuberculosis (Edinb) 2023, 140, 102346.

(48) Kries, H.; Niquille, D. L.; Hilvert, D. A subdomain swap strategy for reengineering nonribosomal peptides. Chem Biol 2015, 22 (5), 640–648.

(49) Baltz, R. H. Combinatorial biosynthesis of cyclic lipopeptide antibiotics: a model for synthetic biology to accelerate the evolution of secondary metabolite biosynthetic pathways. ACS Synth Biol 2014, 3 (10), 748–758.

(50) Baltz, R. H. Synthetic biology, genome mining, and combinatorial biosynthesis of NRPS-derived antibiotics: a perspective. J Ind Microbiol Biotechnol 2018, 45 (7), 635–649.

(51) Folger, I. B.; Frota, N. F.; Pistofidis, A.; Niquille, D. L.; Hansen, D. A.; Schmeing, T. M.; Hilvert, D. High-throughput reprogramming of an NRPS condensation domain. Nat Chem Biol 2024, 20 (6), 761–769.

(52) Peng, H.; Schmiederer, J.; Chen, X.; Panagiotou, G.; Kries, H. Controlling Substrate-and Stereospecificity of Condensation Domains in Nonribosomal Peptide Synthetases. ACS Chem Biol 2024, 19 (3), 599–606.

(53) Adams, P. D.; Afonine, P. V.; Bunkoczi, G.; Chen, V. B.; Davis, I. W.; Echols, N.; Headd, J. J.; Hung, L. W.; Kapral, G. J.; Grosse-Kunstleve, R. W.;, et al. PHENIX: a comprehensive Python-based system for macromolecular structure solution. Acta Crystallogr D Biol Crystallogr 2010, 66 (Pt 2), 213–221.

(54) Emsley, P.; Cowtan, K. Coot: model-building tools for molecular graphics. Acta Crystallogr D Biol Crystallogr 2004, 60 (Pt 12 Pt 1), 2126-2132.

(55) Horswill, A. R.; Escalante-Semerena, J. C. Characterization of the propionyl-CoA synthetase (PrpE) enzyme of Salmonella enterica: residue Lys592 is required for propionyl-AMP synthesis. Biochemistry 2002, 41 (7), 2379–2387.

(56) Wu, M. X.; Hill, K. A. A continuous spectrophotometric assay for the aminoacylation of transfer RNA by alanyl-transfer RNA synthetase. Anal Biochem 1993, 211 (2), 320–323.

