## Supporting Information for "Expanding the substrate selectivity of the fimsbactin biosynthetic adenylation domain, FbsH"

**Syed Fardin Ahmed,[1] Adam Balutowski,[2] Jinping Yang,[2]**

**Timothy A. Wencewicz,[2] Andrew M. Gulick,[1]**

[1]Department of Structural Biology, University at Buffalo, Buffalo, NY, 14203, United States.

[2]Department of Chemistry, Washington University in St. Louis, St. Louis, MO, 63130, United States.

**Supplemental Figures**

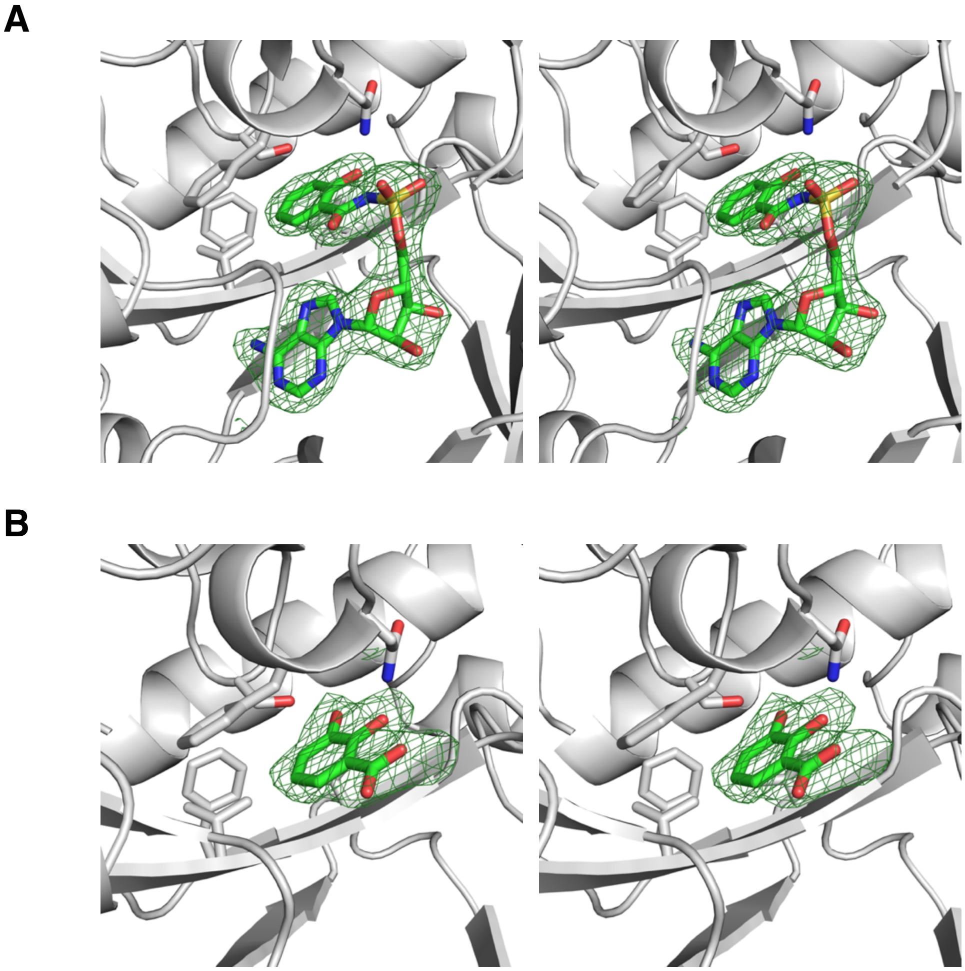

**Supplemental Figure 1**. Density of ligands bound to FbsH. Stereorepresentation of omit maps for the ligands (A) Salicyl-AMS and (B) DHB. The Fo-Fc maps are contoured at 3σ.

```
FbsH       WFRESIVSRS-SIEA---SLPQLQGVAPAFNLRAQSESEDIAFLQLSGGTTGLPKLIPRT    214
3O82       SPEIILMLNHQATDFGLLDWI--ETPAETFVDFSSTPADEVAFFQLSGGSTGTPKLIPRT    210
3RG2       SIRVVQLLND-SGEHNLQDAI--NHPAEDFTA-TPSPADEVAYFQLSGGTTGTPKLIPRT    203
7TYB       C-RMALIHGE-AEAP-LQALAPLYQ--ADALEDCAARAEDIACFQLSGGTTGTPKLIPRR    213
5KEI       DVRHVLVDGD-AAEF--LSWAEVTRAAPGPVPEIAPDPAAPALLLVSGGTTGAPKLIPRT    226
1MD9       TLKNIIVAGE-AEEF--LPLEDLH---TEPVKLPEVKSSDVAFLQLSGGSTGLSKLIPRT    203
5WM2       SIEHVLVAGE-AAEF--TALADVD---AAPVPLAEPDPGDVALLLLSGGTTGKPKLIPRT    204
                      :                             *  : :***:**  *****

FbsH       HADYIYSIEKSVDVAGLTQDTKQLVVLPVMHNFCMSSPGFLGVFYVGGTVVLSQLTHPRV    274
3O82       HNDYDYSVRASAEICGLNSNTRLLCALPAPHNFMLSSPGALGVLHAGGCVVMAPNPEPLN    270
3RG2       HNDYYYSVRRSVEICQFTQQTRYLCAIPAAHNYAMSSPGSLGVFLAGGTVVLAADPSATL    263
7TYB       HREYLYNVRASAEVCGFDEHTVYLTGLPMAHNFTLCCPGVIGTLLASGRVVVSQRADPEH    273
5KEI       HQDYVYNATASAELCRLTADDVYLVALPAAHNFPLACPGLLGAMTVGATTVFTTDPSPEA    286
1MD9       HDDYIYSLKRSVEVCWLDHSTVYLAALPMAHNYPLSSPGVLGVLYAGGRVVLSPSPSPDD    263
5WM2       HDDYTYNVRASAEVCGFDSDTVYLVVLPTAHNFALACPGLLGTLMVGGTVVLAPTPSPED    264
           * :* *.   *.::. :      *  :*  **: :..** :*.: ... .*.:

FbsH       CFELIEKYQIQQVSLVPAIATLWLNAES--LKDYDLSSLQVVQVGGAKLLPSLAEQIIDT    332
3O82       CFSIIQRHQVNMASLVPSAVIMWLEKAA--QYKDQIQSLKLLQVGGASFPESLARQVPEV    328
3RG2       CFPLIEKHQVNVTALVPPAVSLWLQALIEGESRAQLASLKLLQVGGARLSATLAARIPAE    323
7TYB       CFALIARERVTHTALVPPLAMLWLDAQE--SRRADLSSLRLLQVGGSRLGSSAAQRVEPV    331
5KEI       AFAAIDEHGVTATALVPALAKLWAQACA--WEPLAPKTLRLLQVGGAKLAAPDAALVRGA    344
1MD9       AFPLIEREKVTITALVPPLAMVWMDAAS--SRRDDLSSLQVLQVGGAKFSAEAARRVKAV    321
5WM2       AFELIEREKVTATAVVPPVALLWLDAVE--WEDADLSSLRLLQVGGSKLGAEPAARVRPA    322
           .*  * .  :  .::**  . :* :            :*:::****: :    *  :

FbsH       LQVKLQQVYGMAEGLVNFTHLDDSDQITIQTQGKKLSHLDEIRIADQDGNALPINAIGHI    392
3O82       LNCKLQQVFGMAEGLVNYTRLDDSDEQIFTTQGRPISSDDEIKIVDEQYREVPEGEIGML    388
3RG2       IGCQLQQVFGMAEGLVNYTRLDDSAEKIIHTQGYPMCPDDEVWVADAEGNPLPQGEVGRL    383
7TYB       LGCQLQQVLGMAEGLICYTRLDDPPERVLHTQGRPLSPDDEVRVVDAEGREVGPGEVGEL    391
5KEI       LTPGLQQVFGMAEGLLNYTRIGDPPEVLENTQGRPLSPDDEIRIVDEVGNEVPPGAEGEL    404
1MD9       FGCTLQQVFGMAEGLVNYTRLDDPEEIIVNTQGKPMSPYDESRVWDDHDRDVKPGETGHL    381
5WM2       LGCTLQQVFGMAEGLLNYTRLDDPSDLVIQTQGRPLSPDDEIRVVDEDGRDVAPGETGEL    382
           :   **** ******: :*::.*  :    ***  :.  **  : *   . :  .  * :

FbsH       QTRGPYTINGYYNLPEINQRAFTQDGFYKTGDIGYLDEN----LNIVVTGREKEQINRSG    448
3O82       ATRGPYTFCGYYQSPEHNSQVFDEDNYYYSGDLVQRTPD----GNLRVVGRIKDQINRGG    444
3RG2       MTRGPYTFRGYYKSPQHNASAFDANGFYCSGDLISIDPE----GYITVQGREKDQINRGG    439
7TYB       TVRGPYTIRGYYRLPEHNAKAFSADGFYRTGDRVSRDKD----GYLVVEGRDKDQINRGG    447
5KEI       LVRGPYTLNGYFNAEAANERSFSPDGFYRSGDRVRRFADGPLAGYLEVTGRIKDVIVRGG    464
1MD9       LTRGPYTIRGYYKAEEHNAASFTEDGFYRTGDIVRLTRD----GYIVVEGRAKDQINRGG    437
5WM2       LTRGPYTLRGYYRAPEHNARTFSDDGFYRTGDLVRVLPS----GHLVVEGRAKDQINRGG    438
            .*****: **:.    *   *  :.:* :**      .      : * ** *: * *.*
```

**Supplemental Figure 2.** Alignment of the FbsH and adenylating enzyme core sequence. Shown is the core region of FbsH along with BasE (**3O82**), EntE (**3RG2**), PchD (**7TYB**), MbtA (**5KEI**), DhbE (**1MD9**), and CahJ (**5WM2**). The conserved phosphate binding loop and hinge residue between the $A_{core}$ and $A_{sub}$ domains are highlighted in green and red, respectively. The aryl binding pocket is highlighted in yellow, for residues within 5Å of the DHB in FbsH. Phe350 in the second shell is highlighted in cyan.

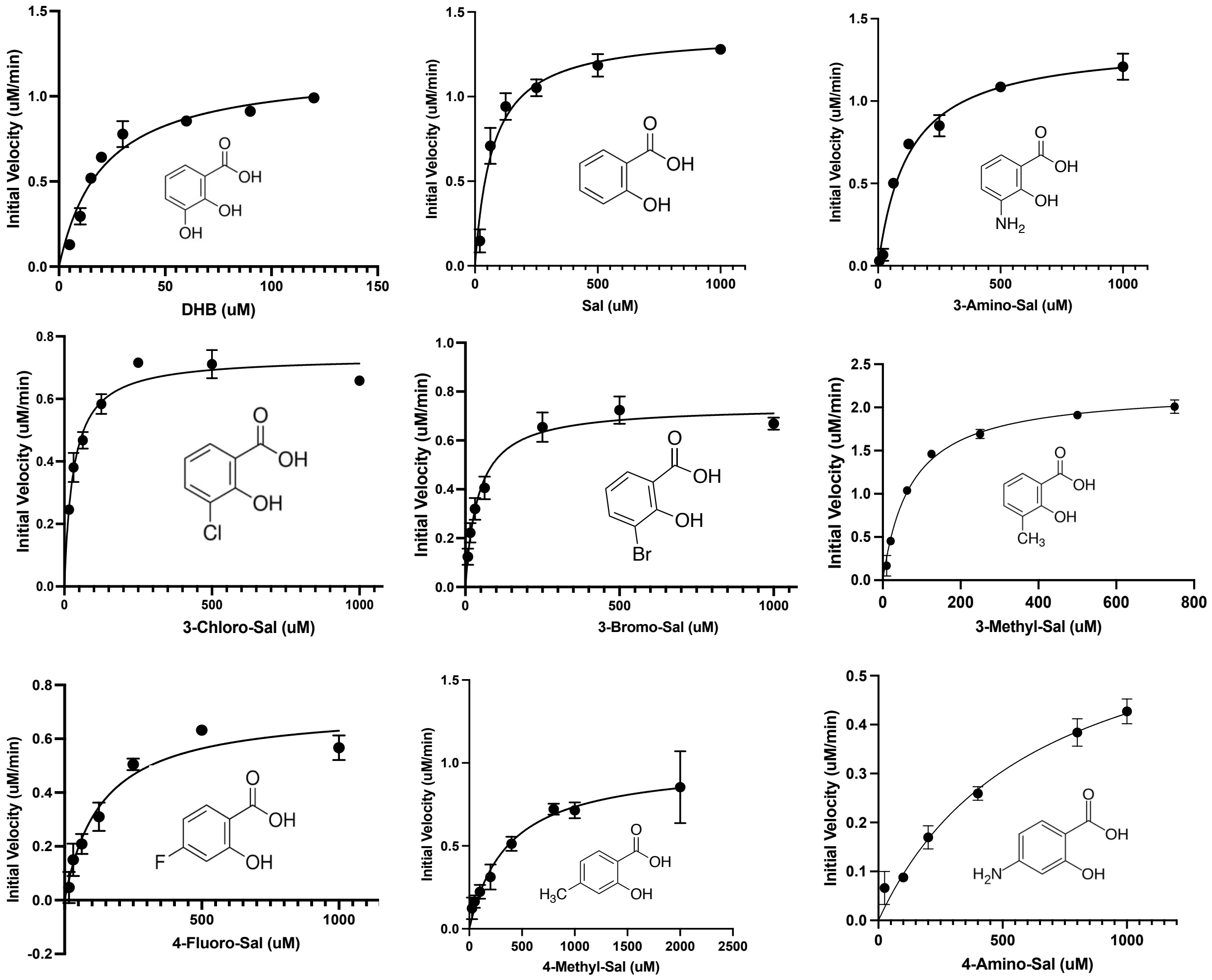

**Supplemental Figure 3**. Michaelis-Menten Kinetics of wildtype FbsH. Initial velocity plots were created for wildtype FbsH against a variety of substrate analogs.

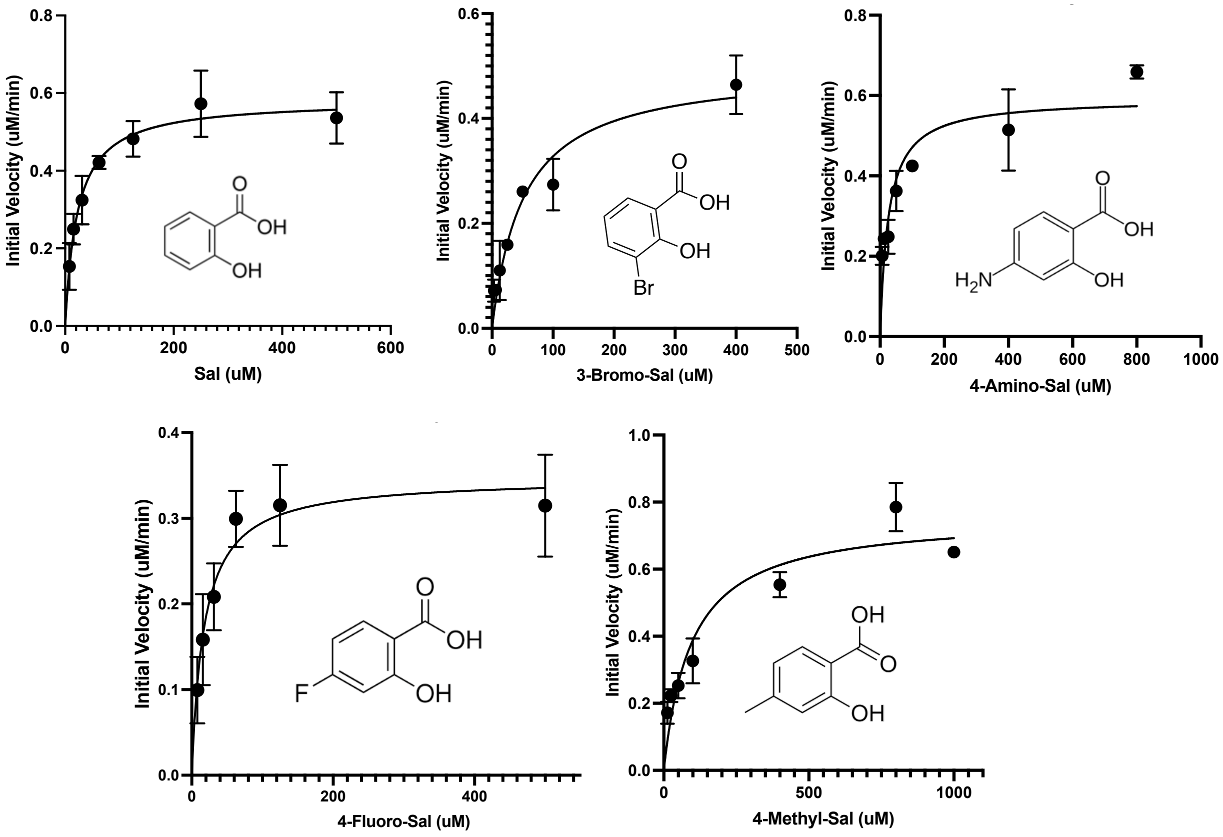

**Supplemental Figure 4**. Michaelis-Menten Kinetics of F350A mutant of FbsH. Initial velocity plots were created for FbsH F350A against a variety of substrate analogs.

**Supplemental Figure 5.** LCMS chromatograms for substrate screen for coupled FbsEFGH enzyme reactions in the presence of FbsM.

Supplemental Figure 5a. 2,3-Dihydroxybenzoic acid

| DHB | FbsH | Aryl-Ser | Aryl-Oxa | Fimsbactin A |
|---|---|---|---|---|
| | Wildtype 1 | 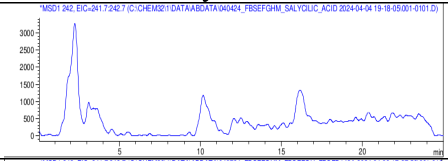 | 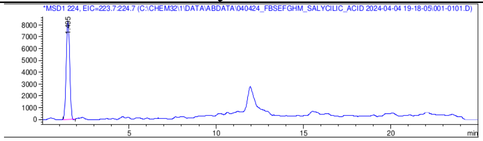 | 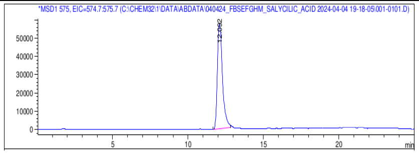 |
| | Wildtype 2 | 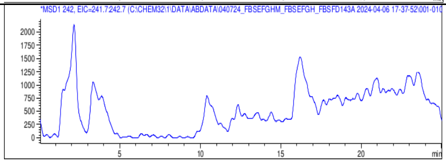 | 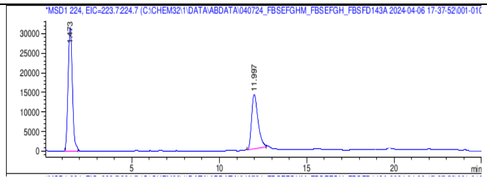 | 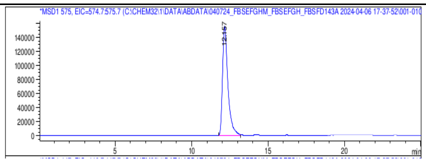 |
| | V348A 1 | 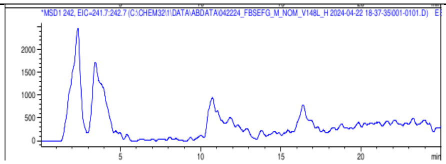 | 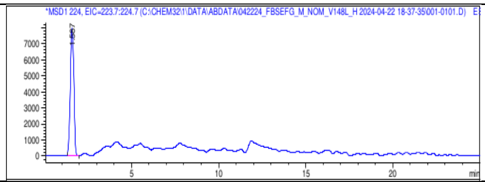 | 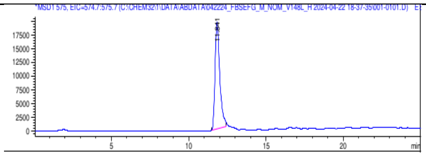 |
| | V348A 2 | 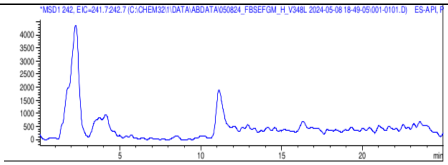 | 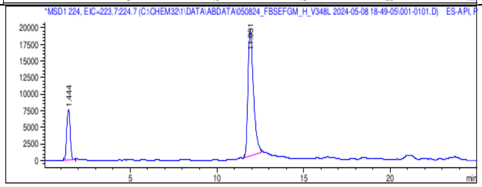 | 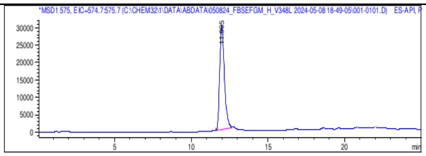 |
| | F350A 1 | 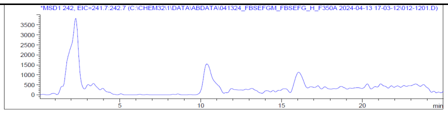 | 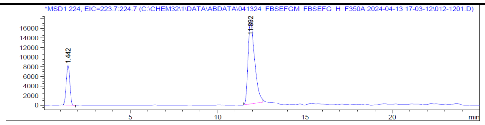 | 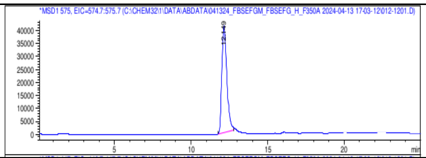 |
| | F350A 2 | 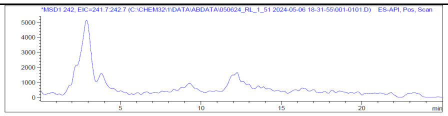 | 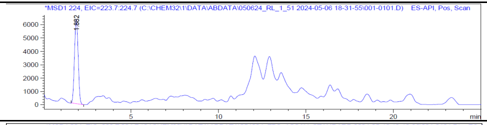 | 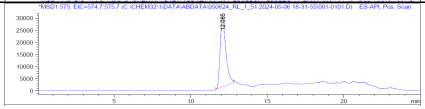 |
| | Standard | 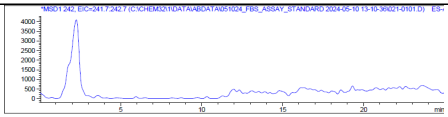 | 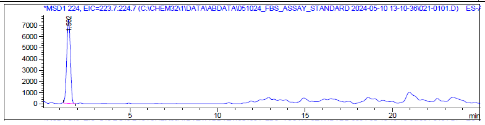 | 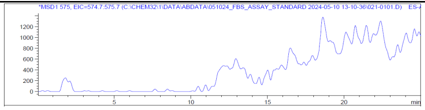 |

Supplemental Figure 5b. Salicylic acid

| | | Aryl-Ser | Aryl-Oxa | Fimsbactin A |
|---|---|---|---|---|
| Sal | FbsH Wildtype 1 | 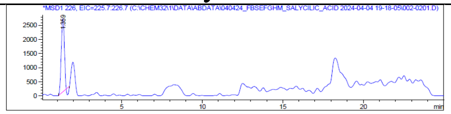 | 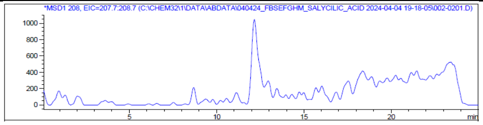 | 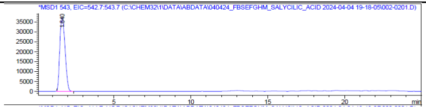 |
| | FbsH Wildtype 2 | 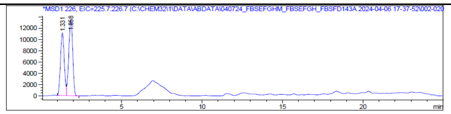 | 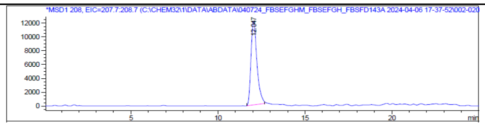 | 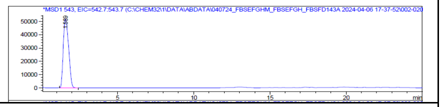 |
| | V348A 1 | 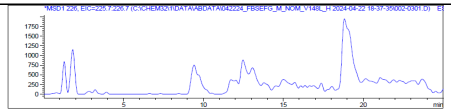 | 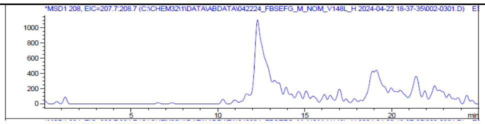 | 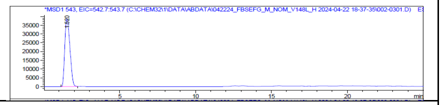 |
| | V348A 2 | 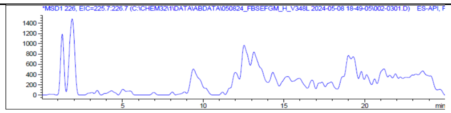 | 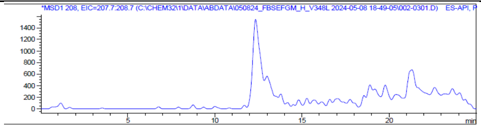 | 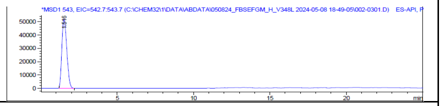 |
| | F350A 1 | 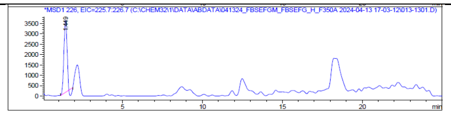 | 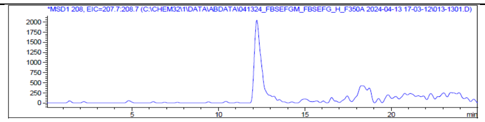 | 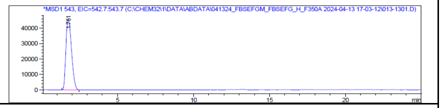 |
| | F350A 2 | 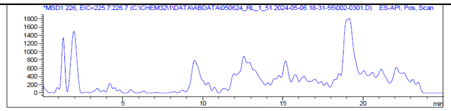 | 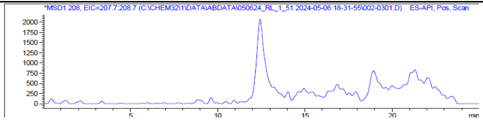 | 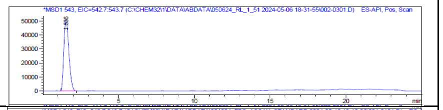 |
| | Standard | 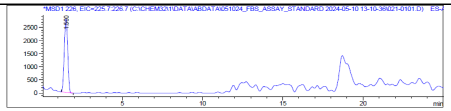 | 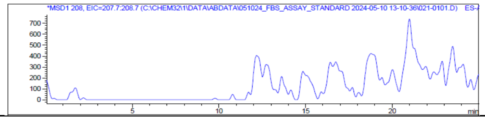 | 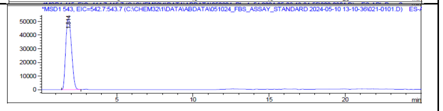 |

Supplemental Figure 5c. 3-Chlorosalicylic acid

| | | Aryl-Ser | Aryl-Oxa | Fimsbactin A |
|---|---|---|---|---|
| 3-Chloro Sal | FbsH Wildtype 1 | 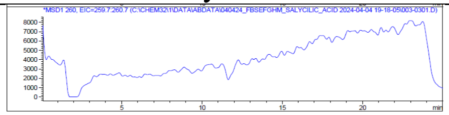 | 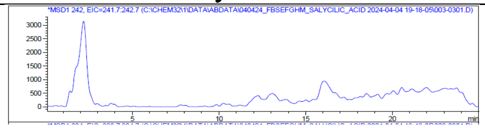 | 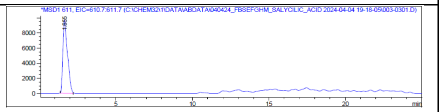 |
| | FbsH Wildtype 2 | 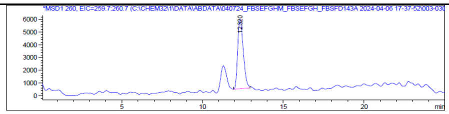 | 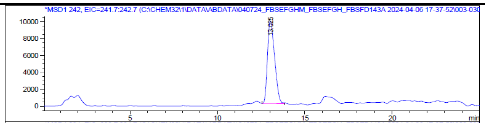 | 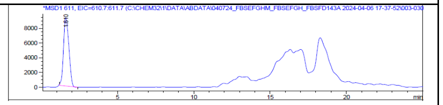 |
| | V348A 1 | 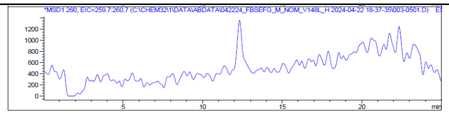 | 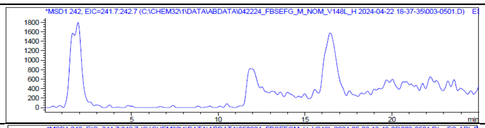 | 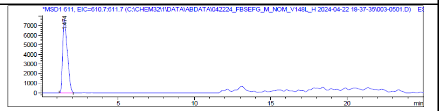 |
| | V348A 2 | 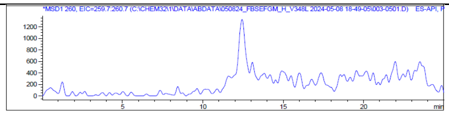 | 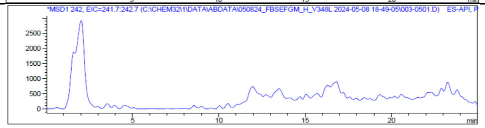 | 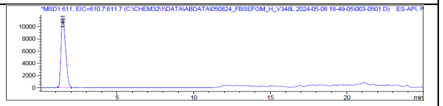 |
| | F350A 1 | 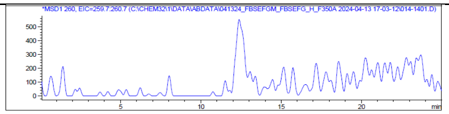 | 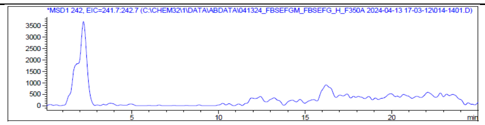 | 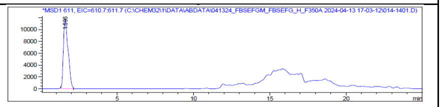 |
| | F350A 2 | 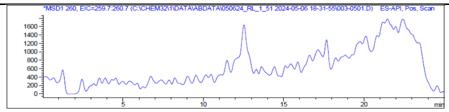 | 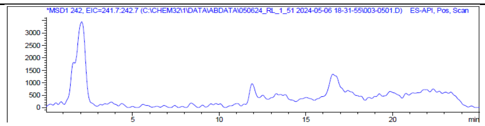 | 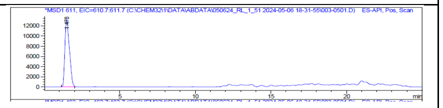 |
| | Standard | 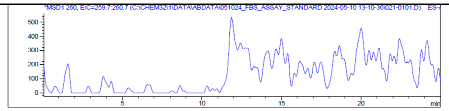 | 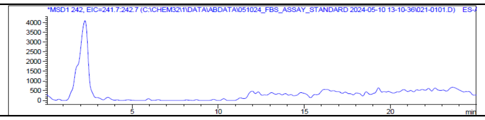 | 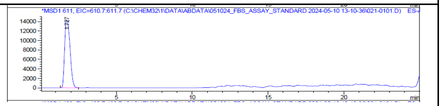 |

Supplemental Figure 5d. 3-Bromosalicylic acid

| | | Aryl-Ser | Aryl-Oxa | Fimsbactin A |
|---|---|---|---|---|
| 3-Bromo Sal | FbsH Wildtype 1 | 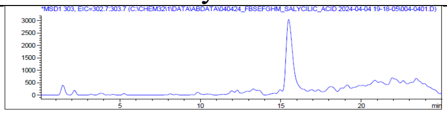 | 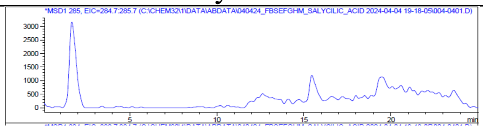 | 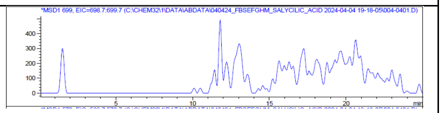 |
| | FbsH Wildtype 2 | 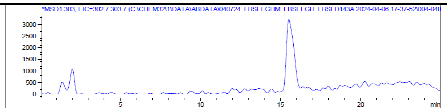 | 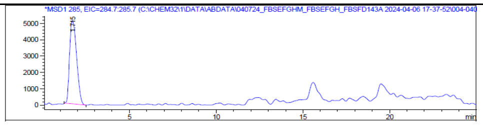 | 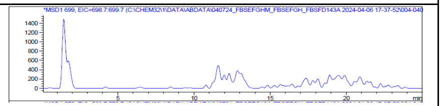 |
| | V348A 1 | 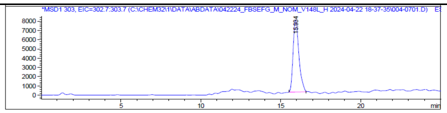 | 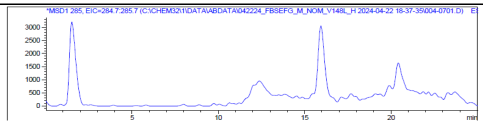 | 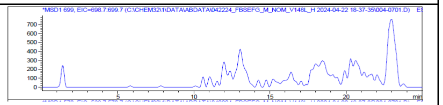 |
| | V348A 2 | 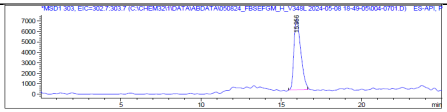 | 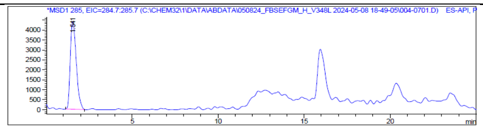 | 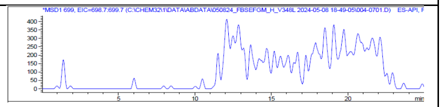 |
| | F350A 1 | 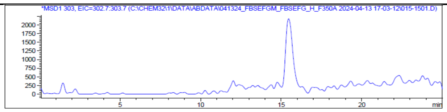 | 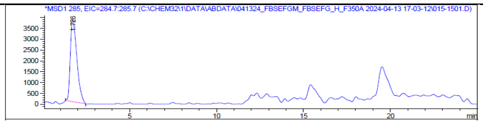 | 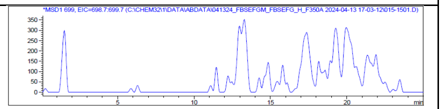 |
| | F350A 2 | 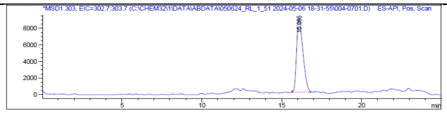 | 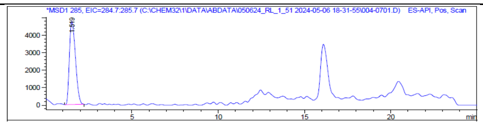 | 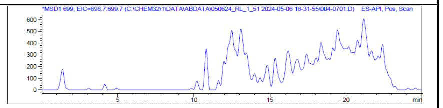 |
| | Standard | 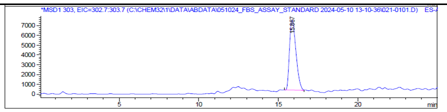 | 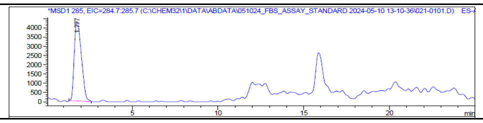 | 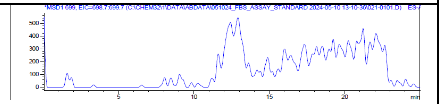 |

Supplemental Figure 5e. 3-Aminosalicylic acid

| | | Aryl-Ser | Aryl-Oxa | Fimsbactin A |
|---|---|---|---|---|
| 3-Amino Sal | FbsH Wildtype 1 | 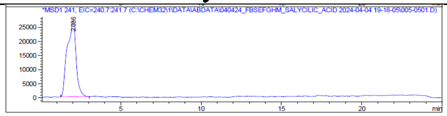 | 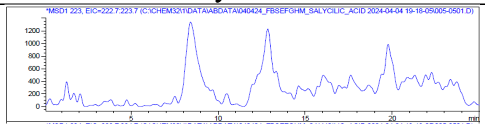 | 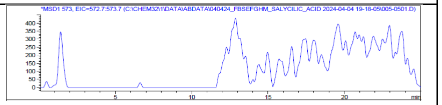 |
| | FbsH Wildtype 2 | 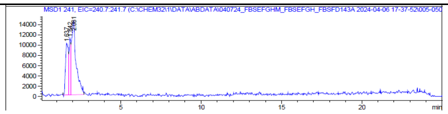 | 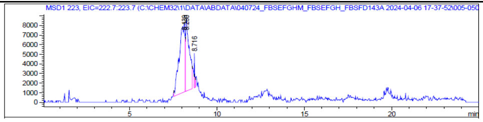 | 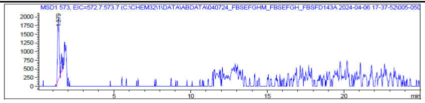 |
| | V348A 1 | 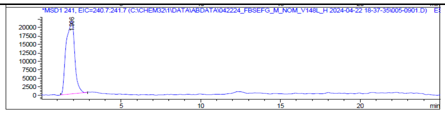 | 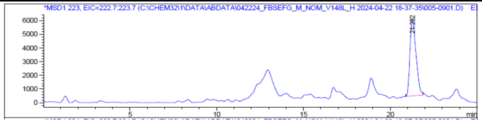 | 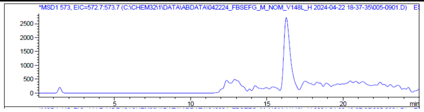 |
| | V348A 2 | 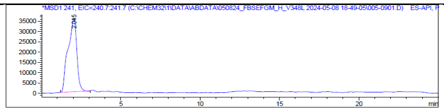 | 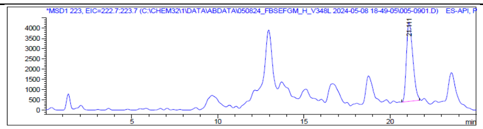 | 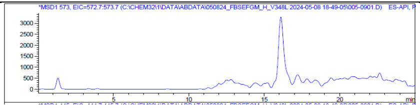 |
| | F350A 1 | 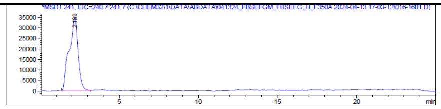 | 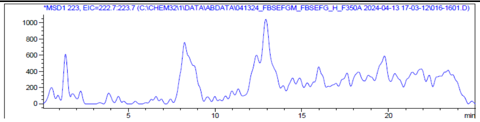 | 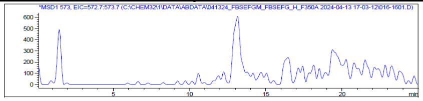 |
| | F350A 2 | 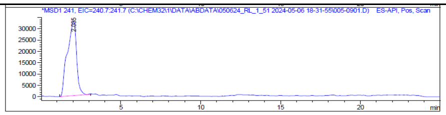 | 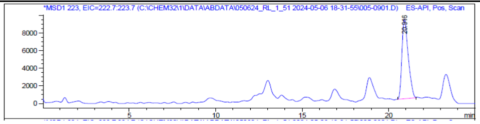 | 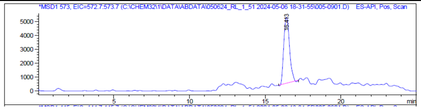 |
| | Standard | 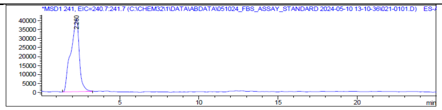 | 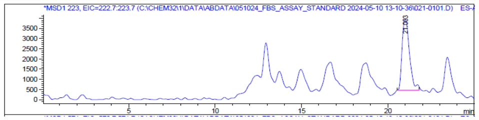 | 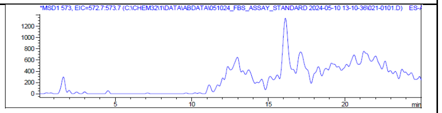 |

Supplemental Figure 5f. 3-Methylsalicylic acid

| | | Aryl-Ser | Aryl-Oxa | Fimsbactin A |
|---|---|---|---|---|
| 3-Methyl Sal | FbsH Wildtype 1 | 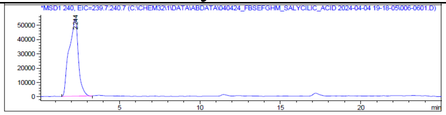 | 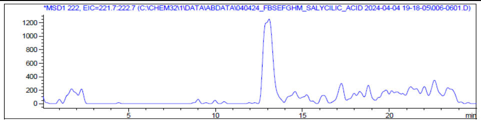 | 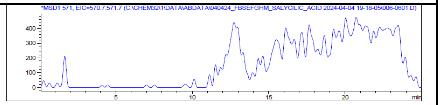 |
| | FbsH Wildtype 2 | 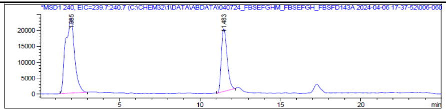 | 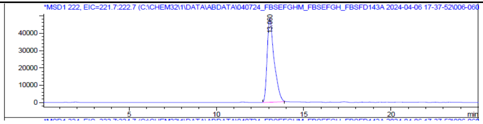 | 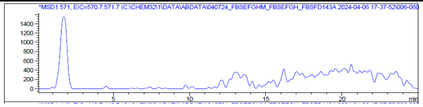 |
| | V348A 1 | 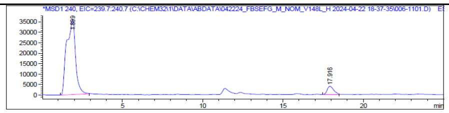 | 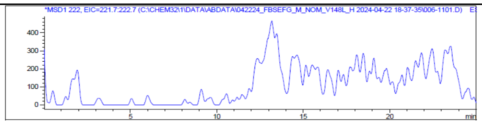 | 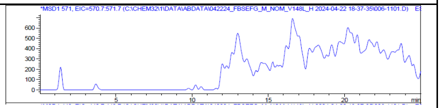 |
| | V348A 2 | 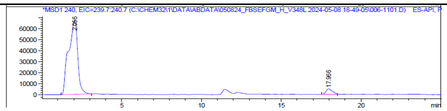 | 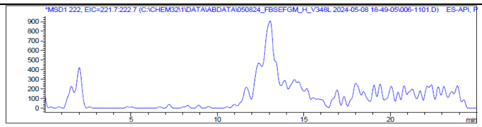 | 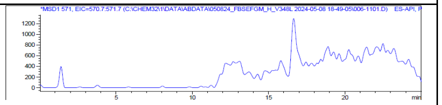 |
| | F350A 1 | 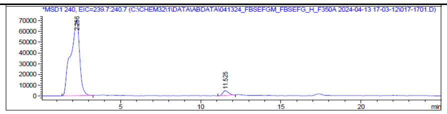 | 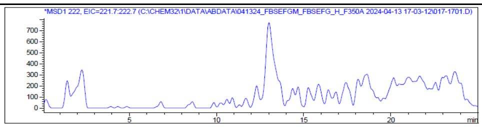 | 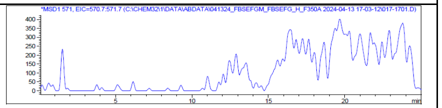 |
| | F350A 2 | 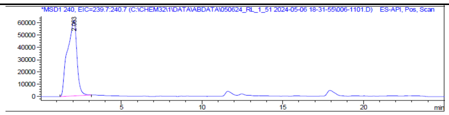 | 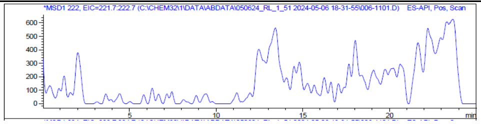 | 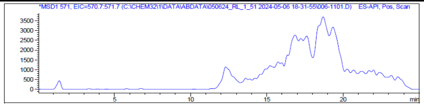 |
| | Standard | 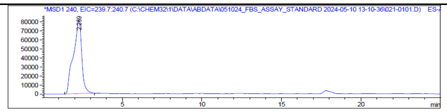 | 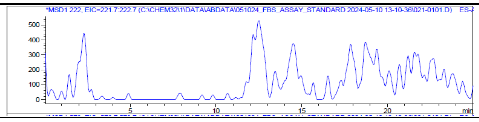 | 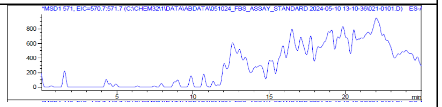 |

Supplemental Figure 5g. 4-Fluorosalicylic acid

| | | Aryl-Ser | Aryl-Oxa | Fimsbactin A |
|---|---|---|---|---|
| 4-Fluoro Sal | FbsH Wildtype 1 | 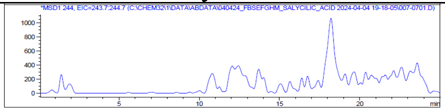 | 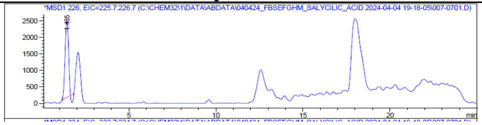 | 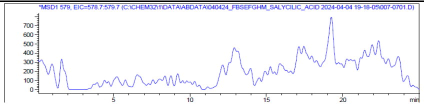 |
| | FbsH Wildtype 2 | 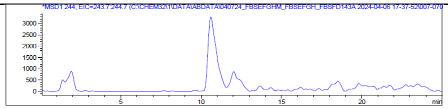 | 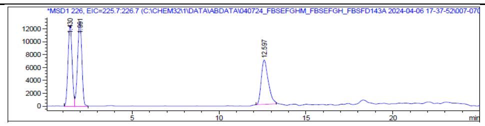 | 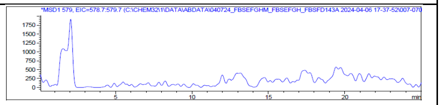 |
| | V348A 1 | 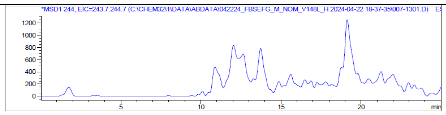 | 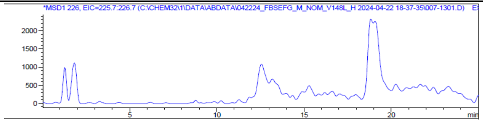 | 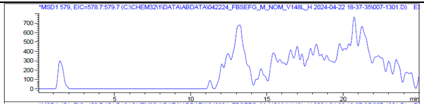 |
| | V348A 2 | 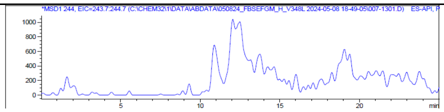 | 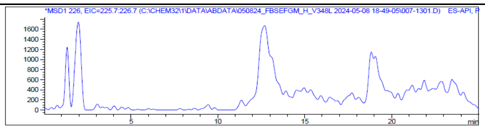 | 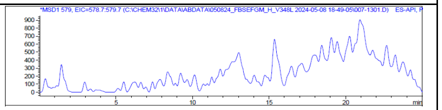 |
| | F350A 1 | 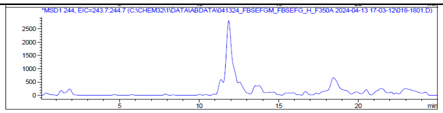 | 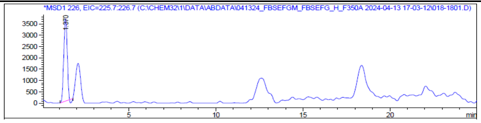 | 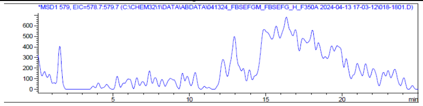 |
| | F350A 2 | 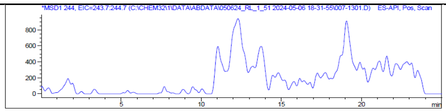 | 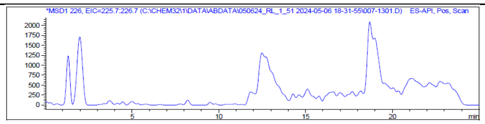 | 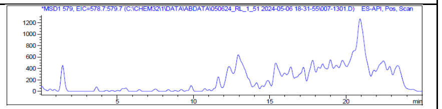 |
| | Standard | 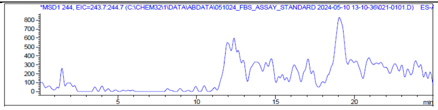 | 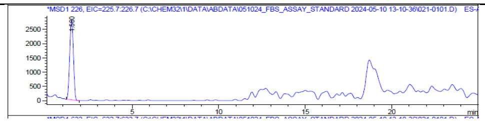 | 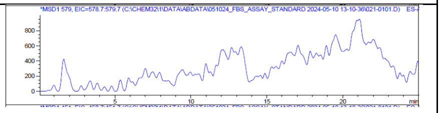 |

Supplemental Figure 5h. 4-Nitrosalicylic acid

| | | Aryl-Ser | Aryl-Oxa | Fimsbactin A |
|---|---|---|---|---|
| 4-Nitro Sal | FbsH Wildtype 1 | 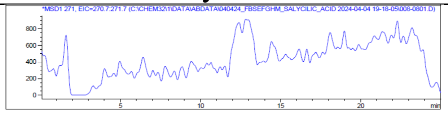 | 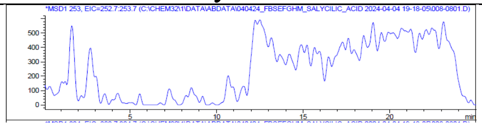 | 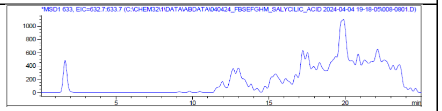 |
| | FbsH Wildtype 2 | 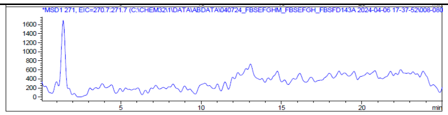 | 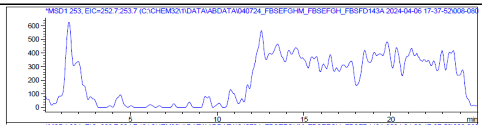 | 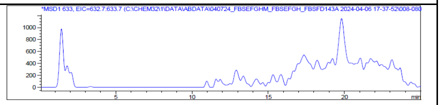 |
| | V348A 1 | 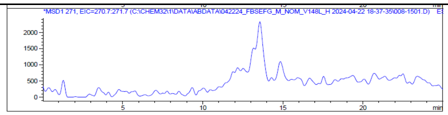 | 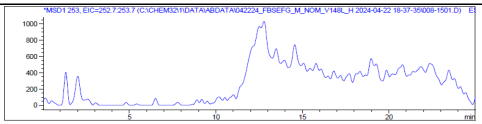 | 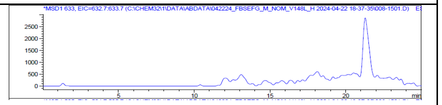 |
| | V348A 2 | 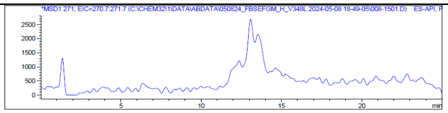 | 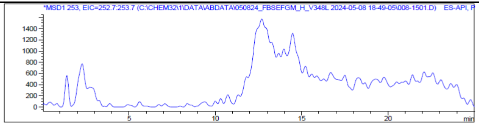 | 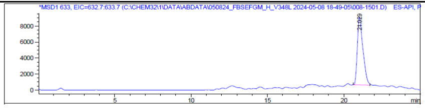 |
| | F350A 1 | 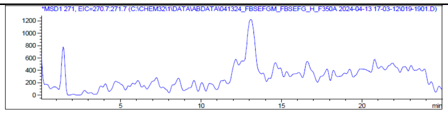 | 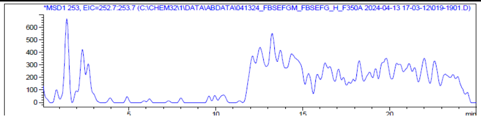 | 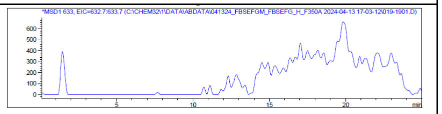 |
| | F350A 2 | 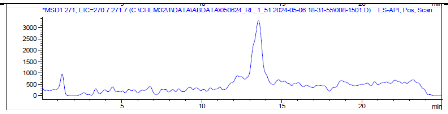 | 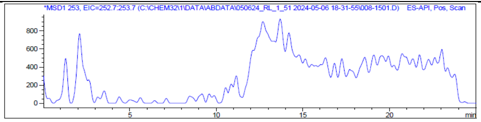 | 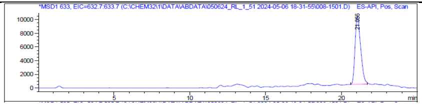 |
| | Standard | 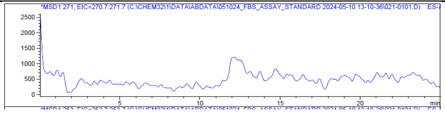 | 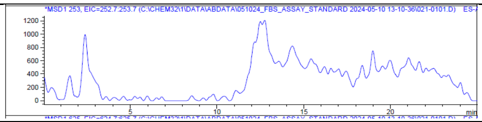 | 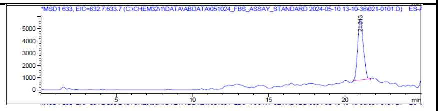 |

Supplemental Figure 5i. 4-Azidosalicylic acid

| | | Aryl-Ser | Aryl-Oxa | Fimsbactin A |
|---|---|---|---|---|
| 4-Azido Sal | FbsH Wildtype 1 | 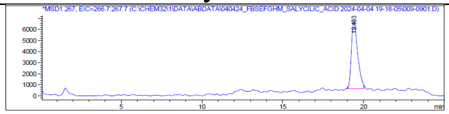 | 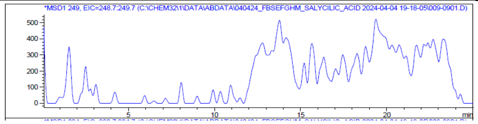 | 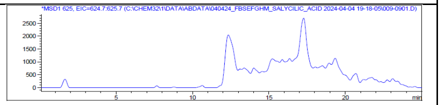 |
| | FbsH Wildtype 2 | 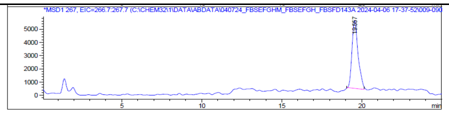 | 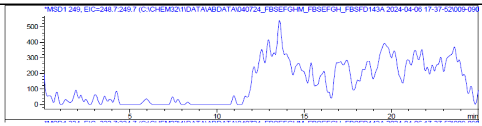 | 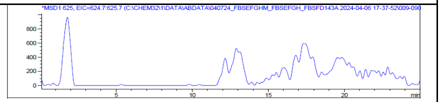 |
| | V348A 1 | 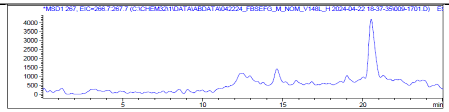 | 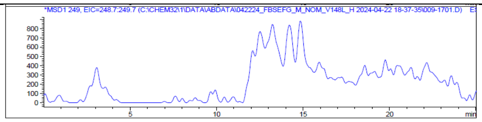 | 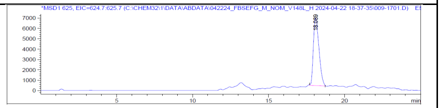 |
| | V348A 2 | 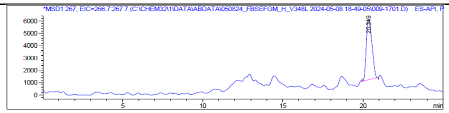 | 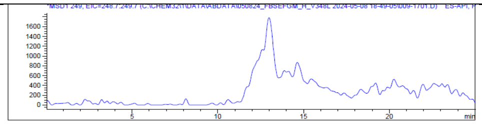 | 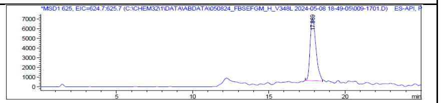 |
| | F350A 1 | 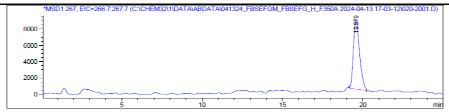 | 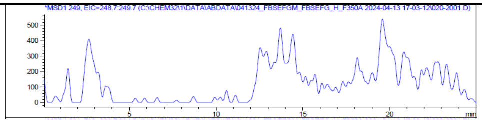 | 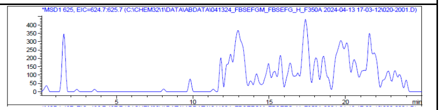 |
| | F350A 2 | 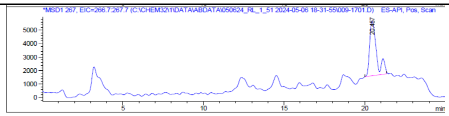 | 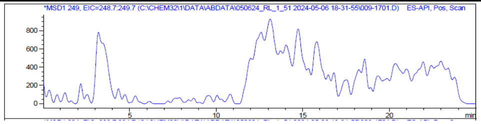 | 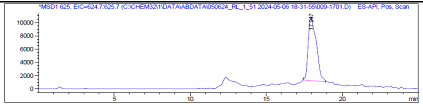 |
| | Standard | 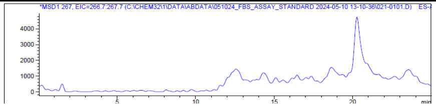 | 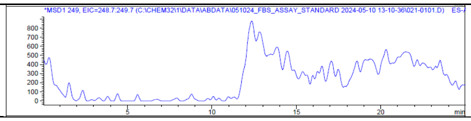 | 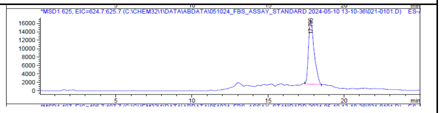 |

Supplemental Figure 5j. 4-Aminosalicylic acid

| | | Aryl-Ser | Aryl-Oxa | Fimsbactin A |
|---|---|---|---|---|
| 4-Amino Sal | FbsH Wildtype 1 | 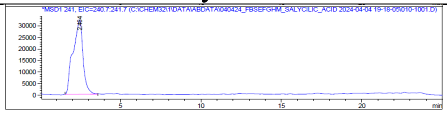 | 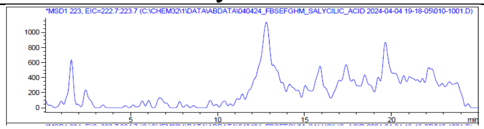 | 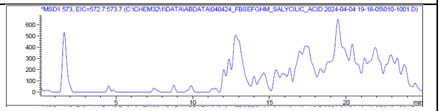 |
| | FbsH Wildtype 2 | 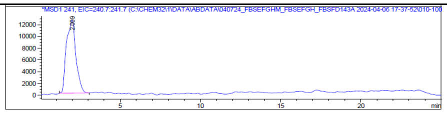 | 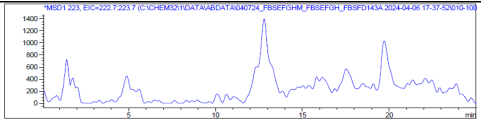 | 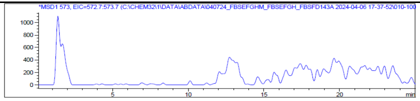 |
| | V348A 1 | 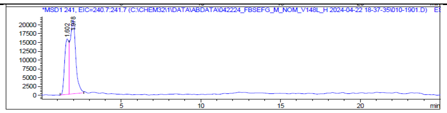 | 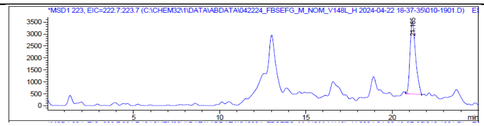 | 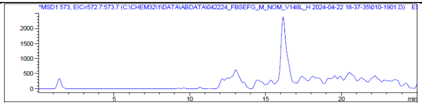 |
| | V348A 2 | 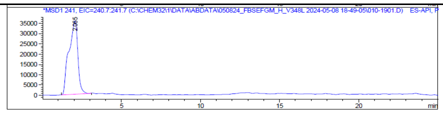 | 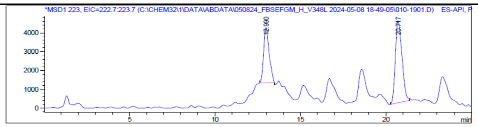 | 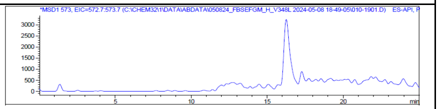 |
| | F350A 1 | 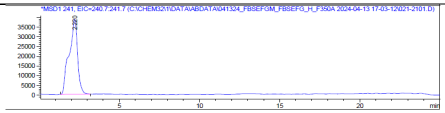 | 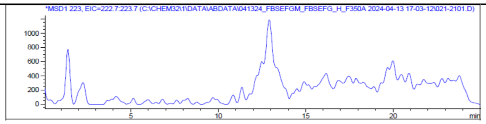 | 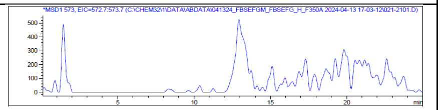 |
| | F350A 2 | 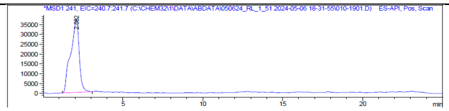 | 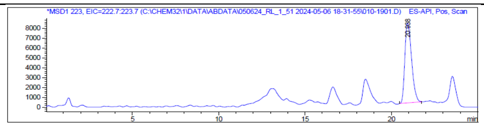 | 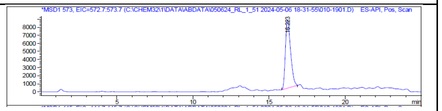 |
| | Standard | 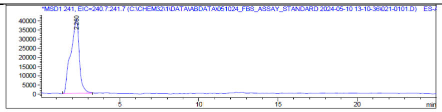 | 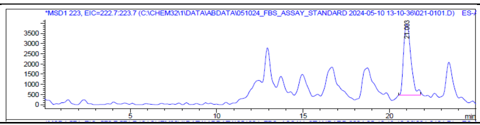 | 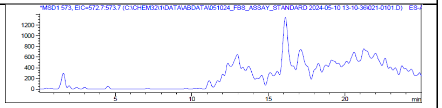 |

Supplemental Figure 5k. 4-Methylsalicylic acid

| | | Aryl-Ser | Aryl-Oxa | Fimsbactin A |
|---|---|---|---|---|
| 4-Methyl Sal | FbsH Wildtype 1 | 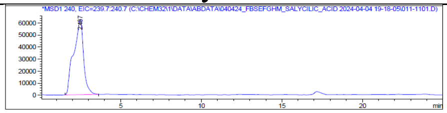 | 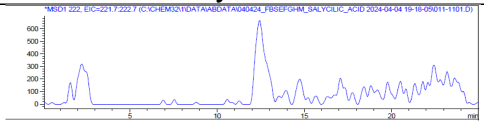 | 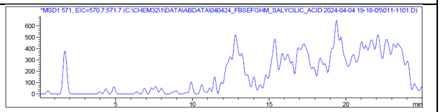 |
| | FbsH Wildtype 2 | 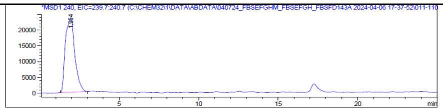 | 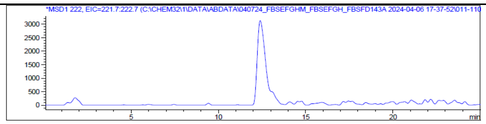 | 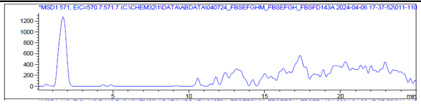 |
| | V348A 1 | 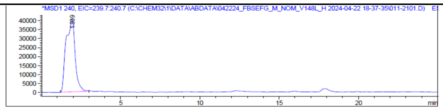 | 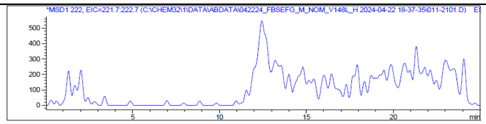 | 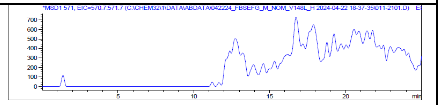 |
| | V348A 2 | 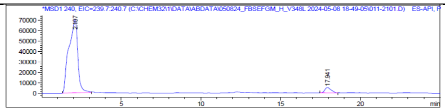 | 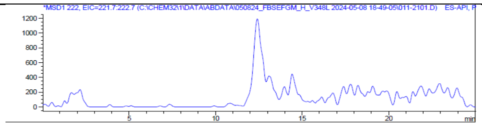 | 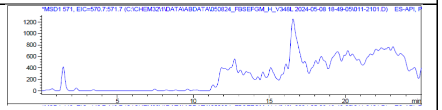 |
| | F350A 1 | 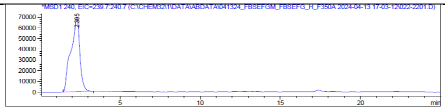 | 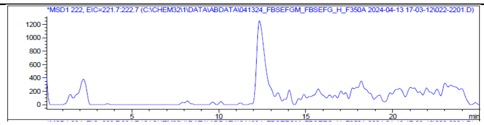 | 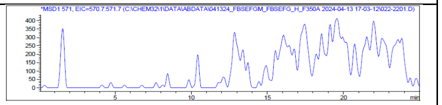 |
| | F350A 2 | 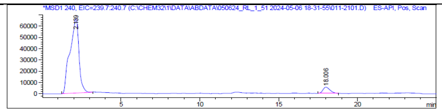 | 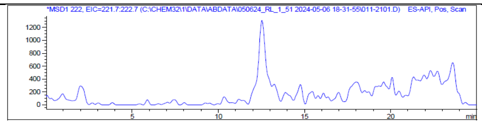 | 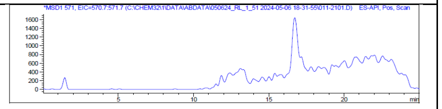 |
| | Standard | 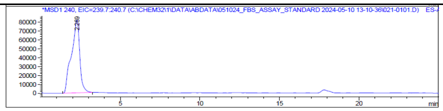 | 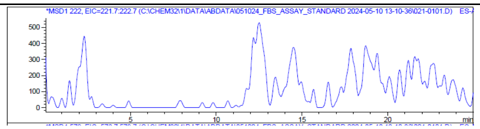 | 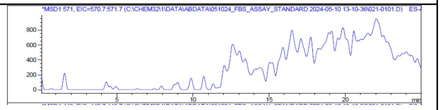 |

**Supplemental Figure 6.** LCMS chromatograms for DHB analogs substrate screen for coupled FbsEFGH enzyme reactions in the absence of FbsM thioesterase.

Supplemental Figure 6a. 2,3-Dihydroxybenzoic acid

| | | Aryl-Ser | Aryl-Oxa | Fimsbactin A |
|---|---|---|---|---|
| DHB | FbsH Wildtype 1 | 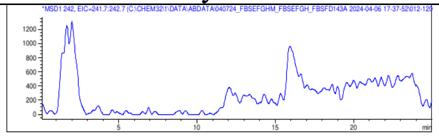 | 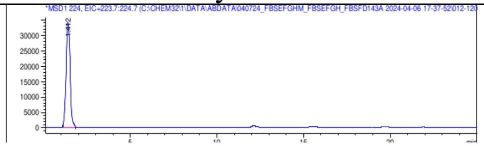 | 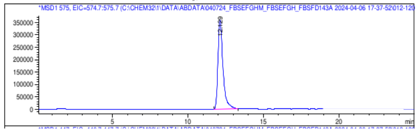 |
| | FbsH Wildtype 2 | 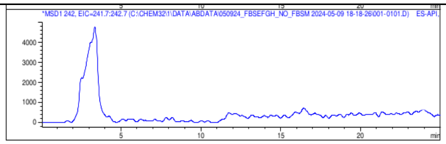 | 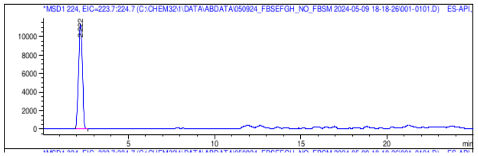 | 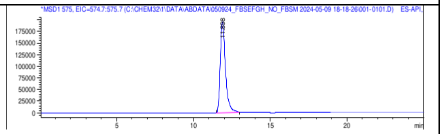 |
| | V348A 1 | 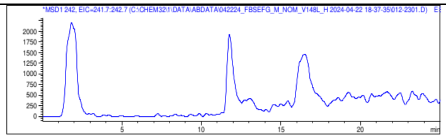 | 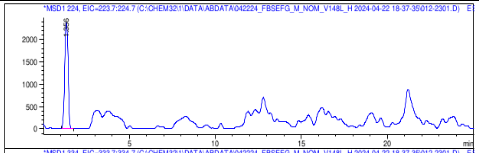 | 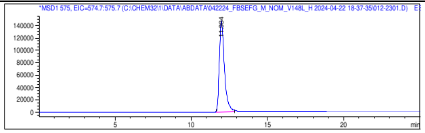 |
| | V348A 2 | 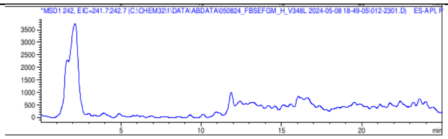 | 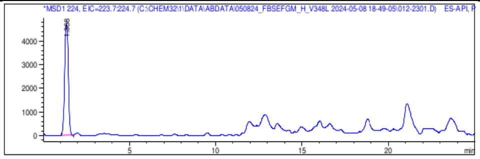 | 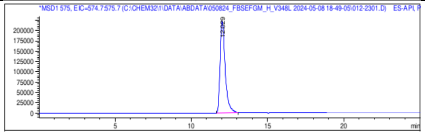 |
| | F350A 1 | 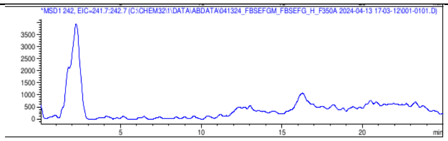 | 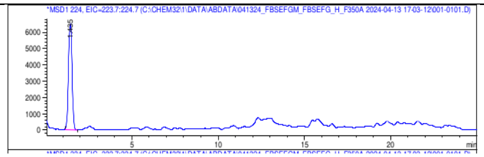 | 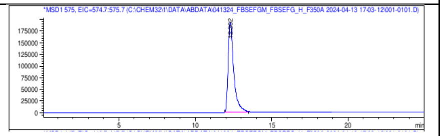 |
| | F350A 2 | 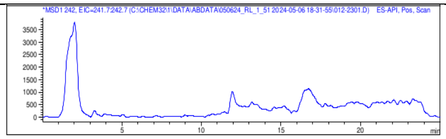 | 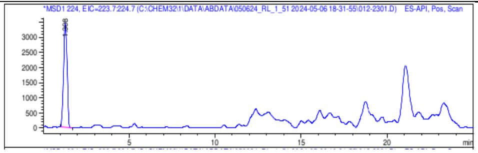 | 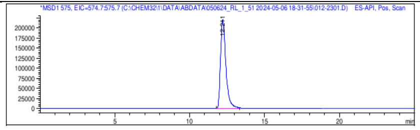 |
| | Standard | 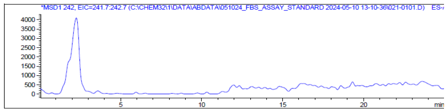 | 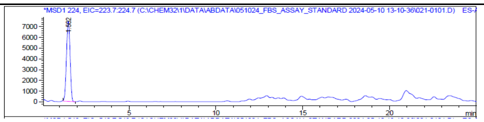 | 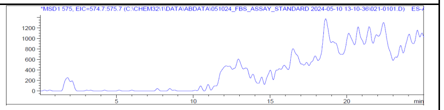 |

Supplemental Figure 6b. Salicylic acid

| | | Aryl-Ser | Aryl-Oxa | Fimsbactin A |
|---|---|---|---|---|
| Sal | FbsH Wildtype 1 | 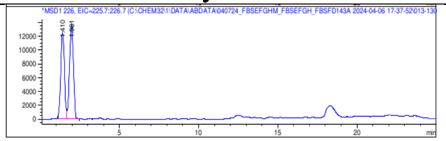 | 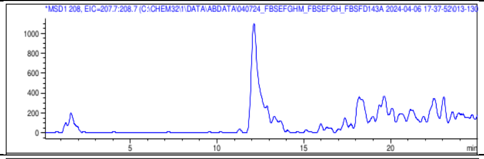 | 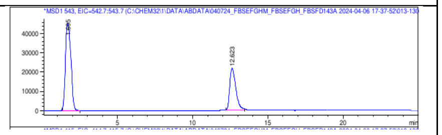 |
| | FbsH Wildtype 2 | 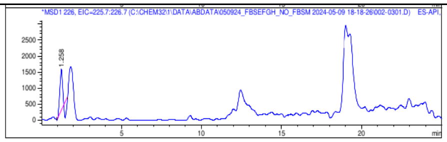 | 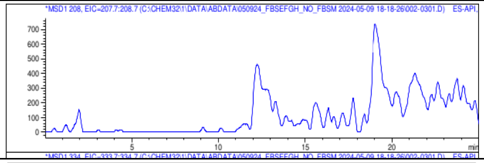 | 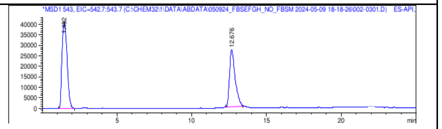 |
| | V348A 1 | 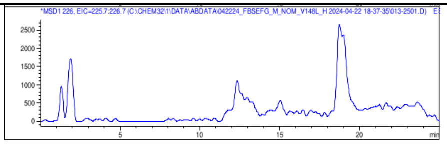 | 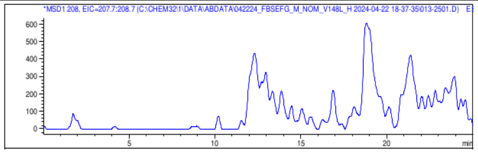 | 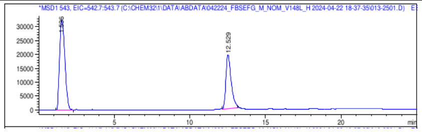 |
| | V348A 2 | 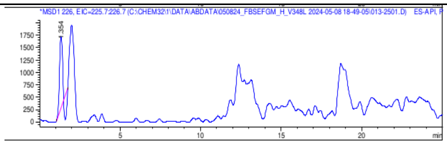 | 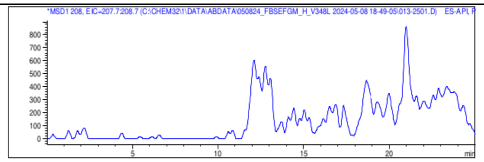 | 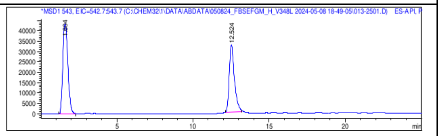 |
| | F350A 1 | 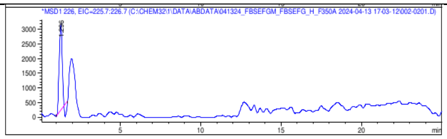 | 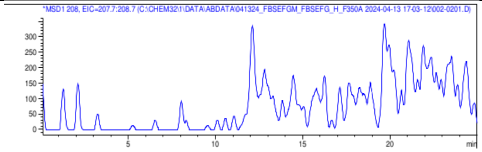 | 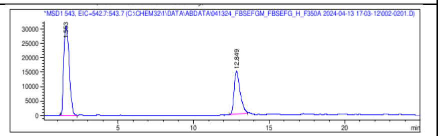 |
| | F350A 2 | 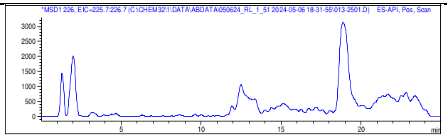 | 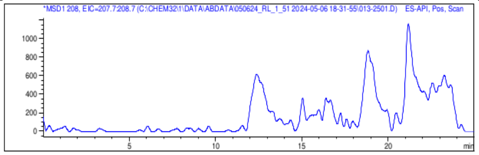 | 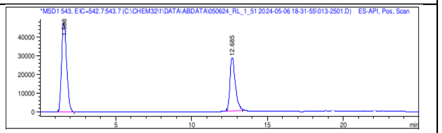 |
| | Standard | 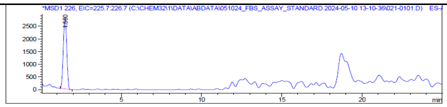 | 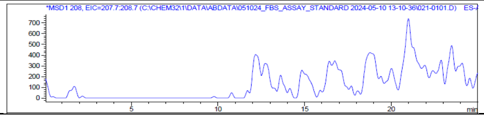 | 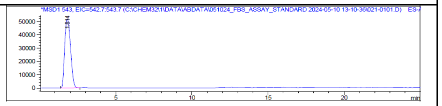 |

Supplemental Figure 6c. 3-Chlorosalicylic acid

| | | Aryl-Ser | Aryl-Oxa | Fimsbactin A |
|---|---|---|---|---|
| 3-Chloro Sal | FbsH Wildtype 1 | 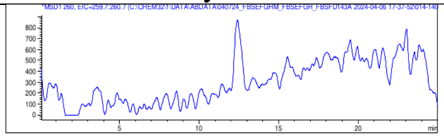 | 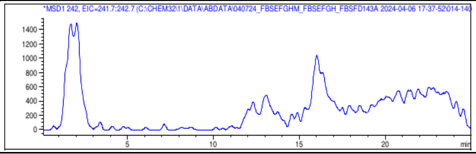 | 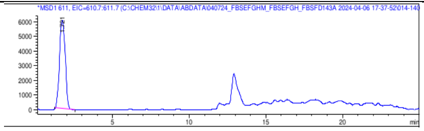 |
| | FbsH Wildtype 2 | 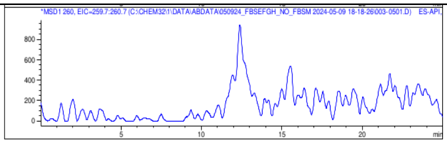 | 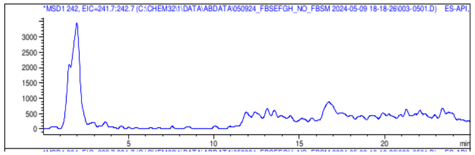 | 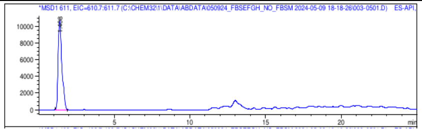 |
| | V348A 1 | 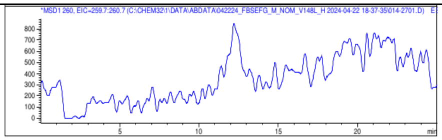 | 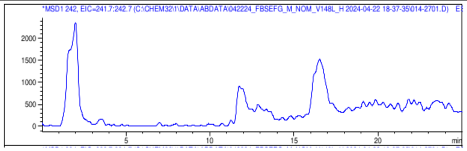 | 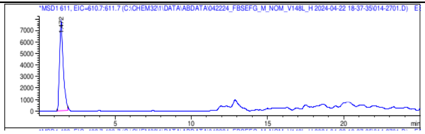 |
| | V348A 2 | 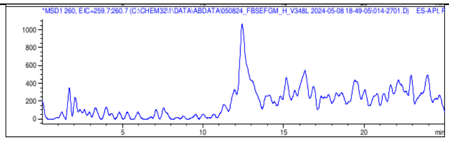 | 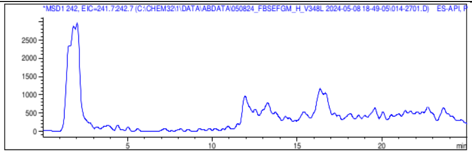 | 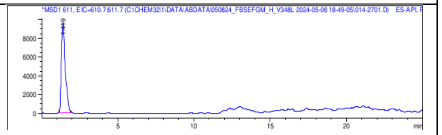 |
| | F350A 1 | 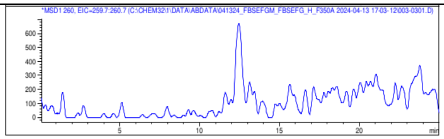 | 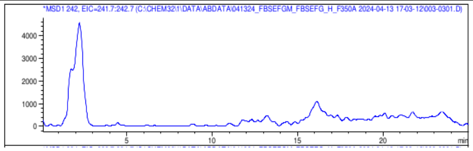 | 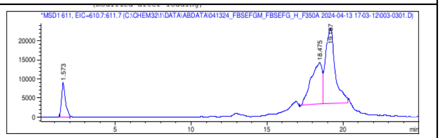 |
| | F350A 2 | 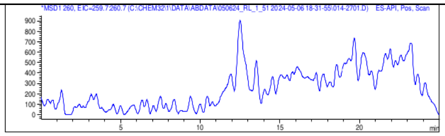 | 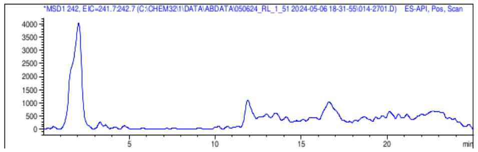 | 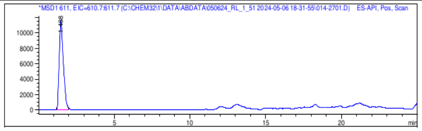 |
| | Standard | 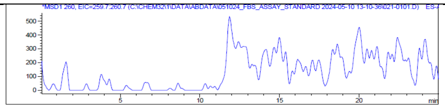 | 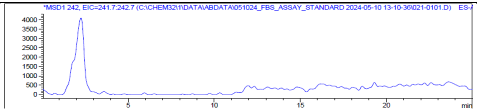 | 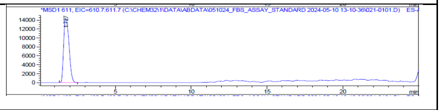 |

Supplemental Figure 6d. 3-Bromosalicylic acid

| | | Aryl-Ser | Aryl-Oxa | Fimsbactin A |
|---|---|---|---|---|
| 3-Bromo Sal | FbsH Wildtype 1 | 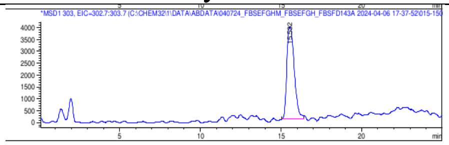 | 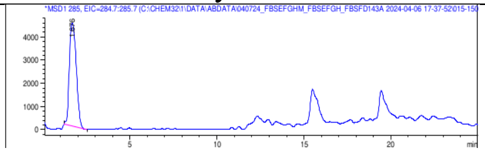 | 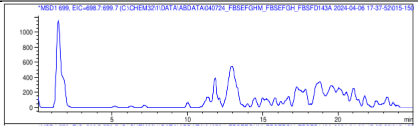 |
| | FbsH Wildtype 2 | 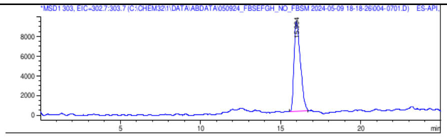 | 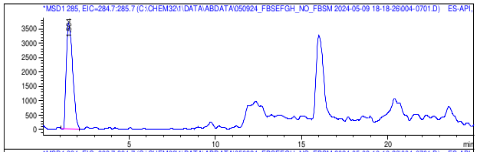 | 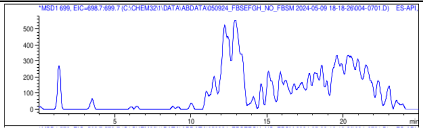 |
| | V348A 1 | 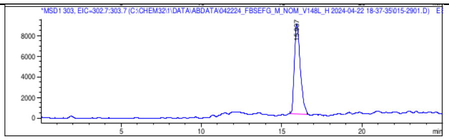 | 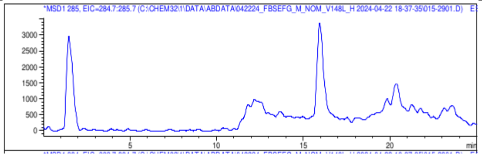 | 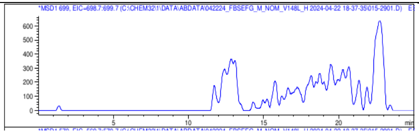 |
| | V348A 2 | 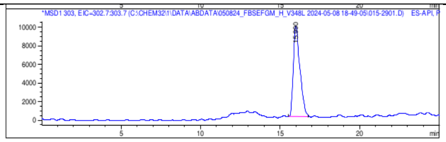 | 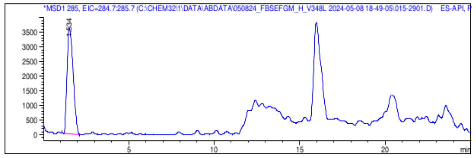 | 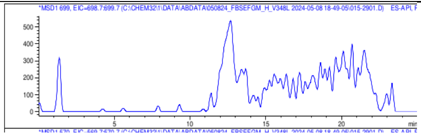 |
| | F350A 1 | 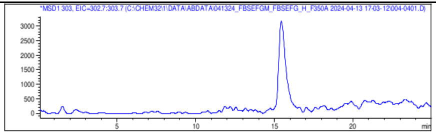 | 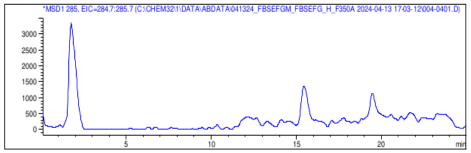 | 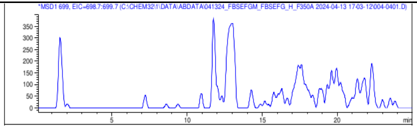 |
| | F350A 2 | 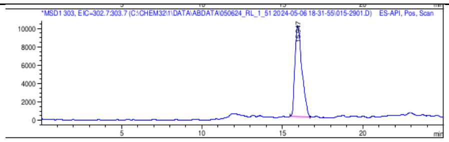 | 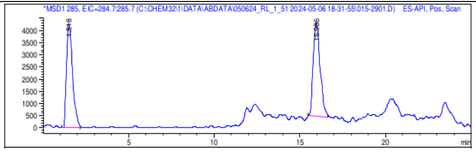 | 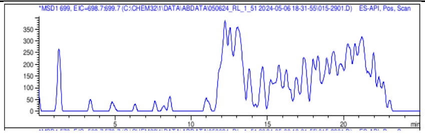 |
| | Standard | 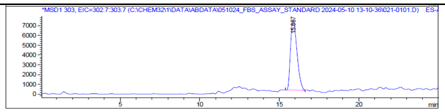 | 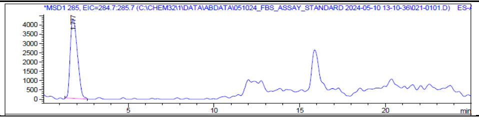 | 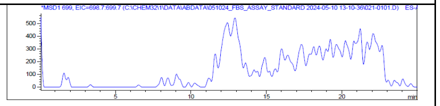 |

Supplemental Figure 6e. 3-Aminosalicylic acid

| | | Aryl-Ser | Aryl-Oxa | Fimsbactin A |
|---|---|---|---|---|
| 3-Amino Sal | FbsH Wildtype 1 | 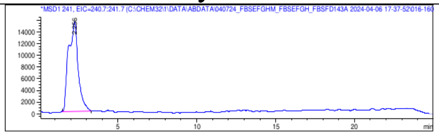 | 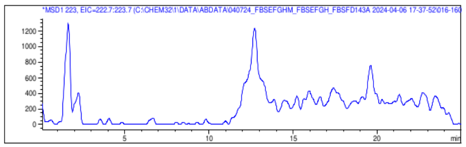 | 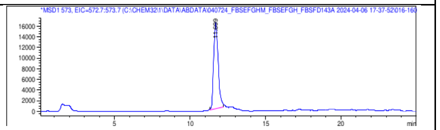 |
| | FbsH Wildtype 2 | 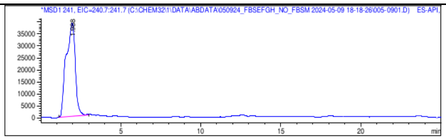 | 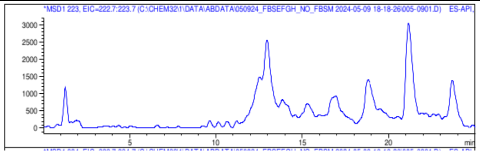 | 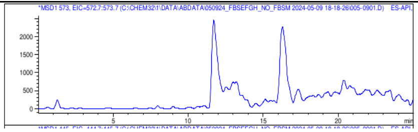 |
| | V348A 1 | 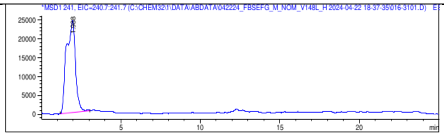 | 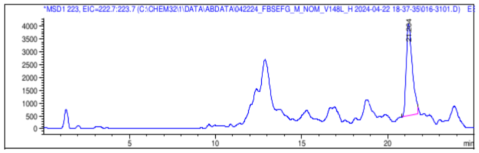 | 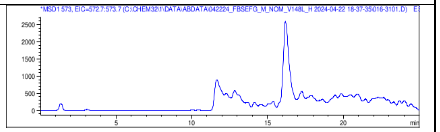 |
| | V348A 2 | 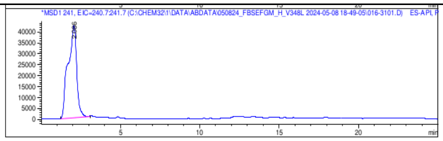 | 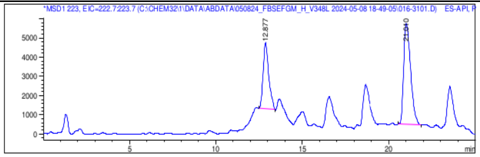 | 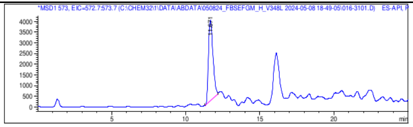 |
| | F350A 1 | 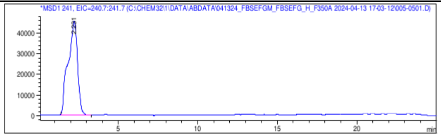 | 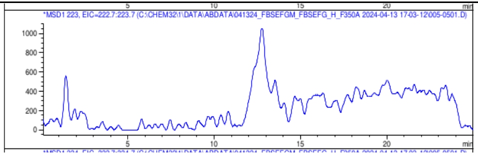 | 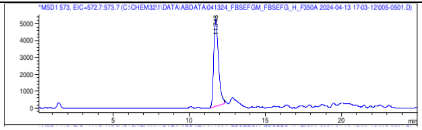 |
| | F350A 2 | 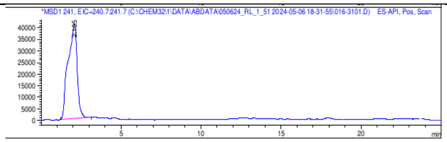 | 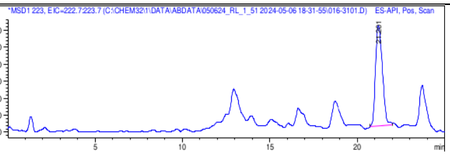 | 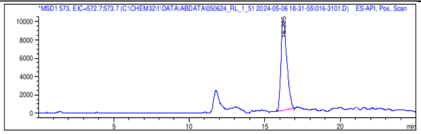 |
| | Standard | 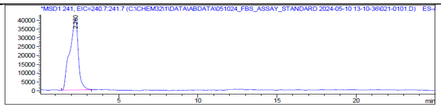 | 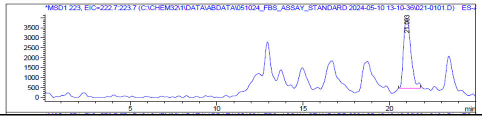 | 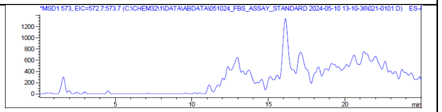 |

Supplemental Figure 6f. 3-Methylsalicylic acid

| | | Aryl-Ser | Aryl-Oxa | Fimsbactin A |
|---|---|---|---|---|
| 3-Methyl Sal | FbsH Wildtype 1 | 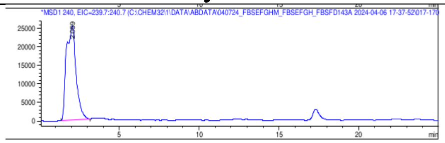 | 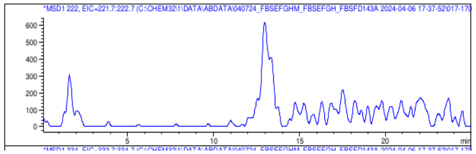 | 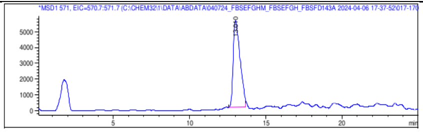 |
| | FbsH Wildtype 2 | 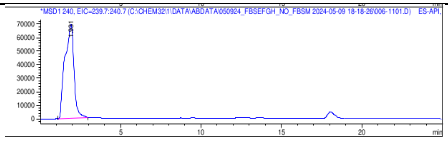 | 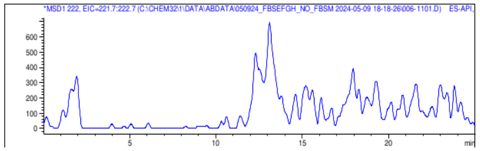 | 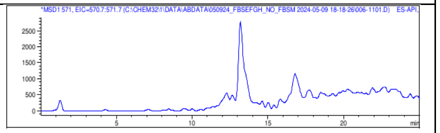 |
| | V348A 1 | 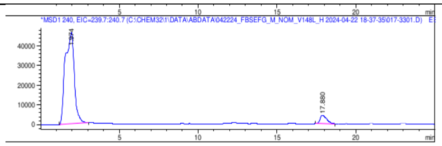 | 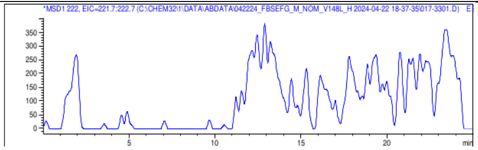 | 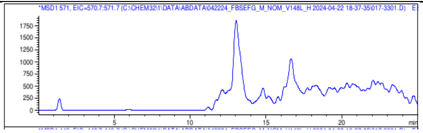 |
| | V348A 2 | 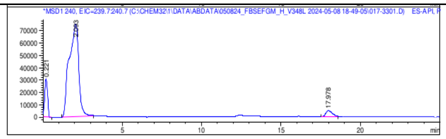 | 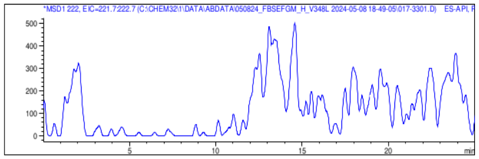 | 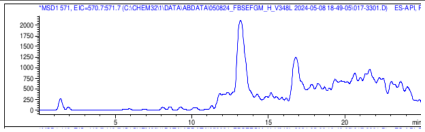 |
| | F350A 1 | 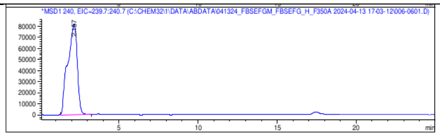 | 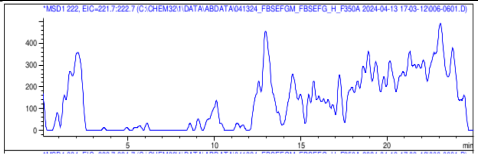 | 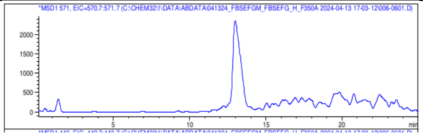 |
| | F350A 2 | 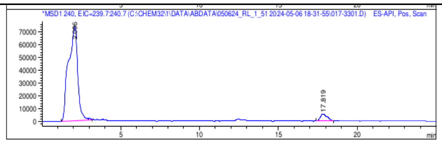 | 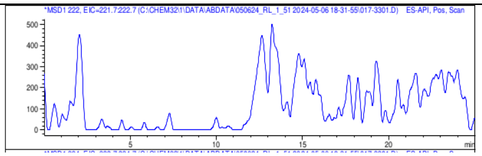 | 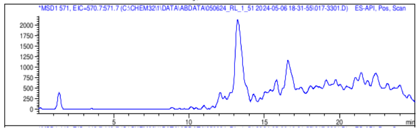 |
| | Standard | 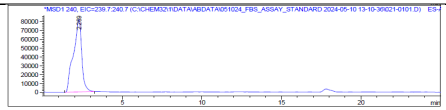 | 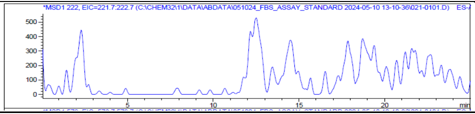 | 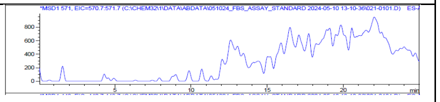 |

Supplemental Figure 6g. 4-Fluorosalicylic acid

| | | Aryl-Ser | Aryl-Oxa | Fimsbactin A |
|---|---|---|---|---|
| 4-Fluoro Sal | FbsH Wildtype 1 | 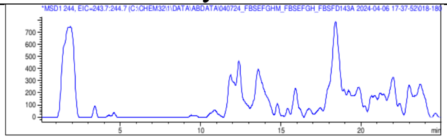 | 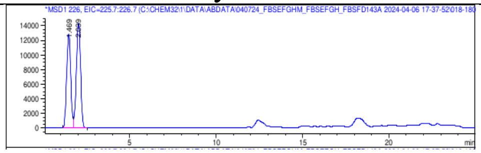 | 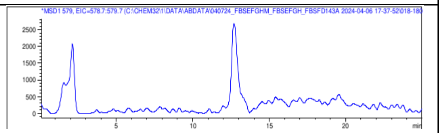 |
| | FbsH Wildtype 2 | 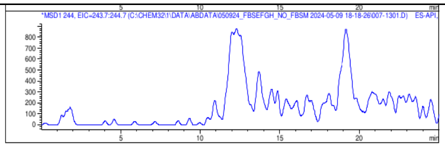 | 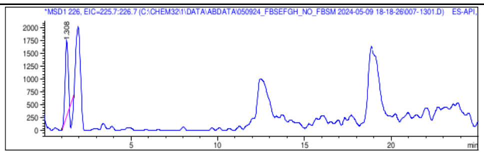 | 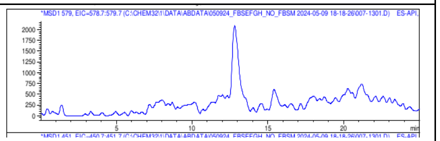 |
| | V348A 1 | 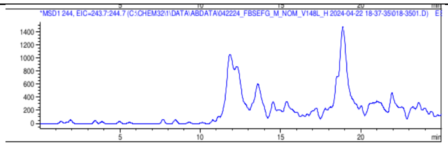 | 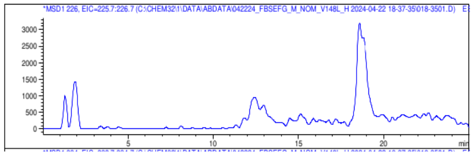 | 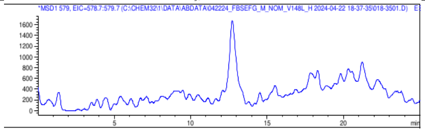 |
| | V348A 2 | 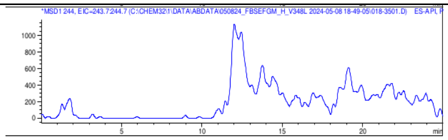 | 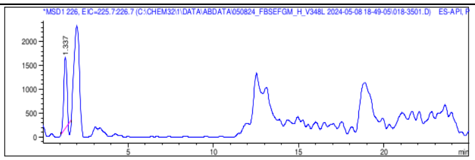 | 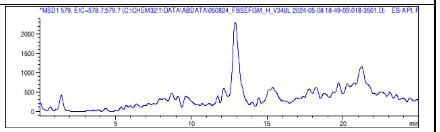 |
| | F350A 1 | 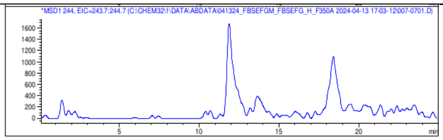 | 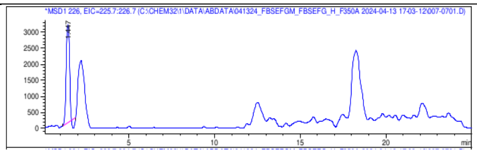 | 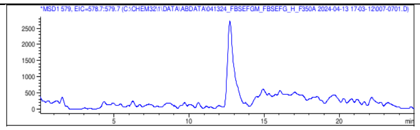 |
| | F350A 2 | 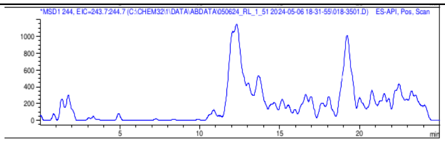 | 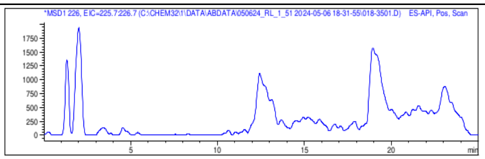 | 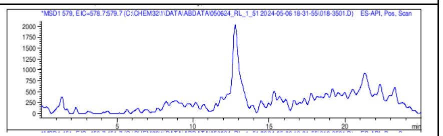 |
| | Standard | 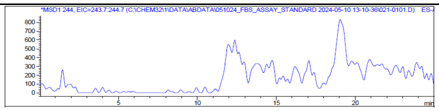 | 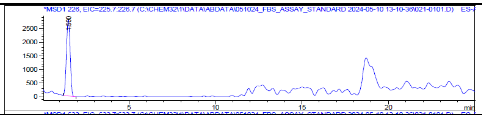 | 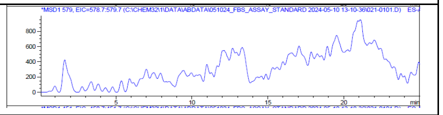 |

Supplemental Figure 6h. 4-Nitrosalicylic acid

| | | Aryl-Ser | Aryl-Oxa | Fimsbactin A |
|---|---|---|---|---|
| 4-Nitro Sal | FbsH Wildtype 1 | 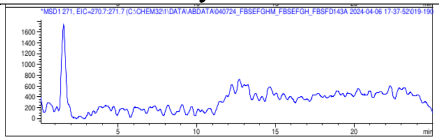 | 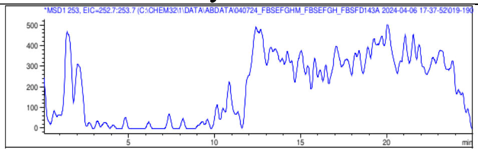 | 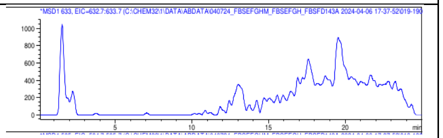 |
| | FbsH Wildtype 2 | 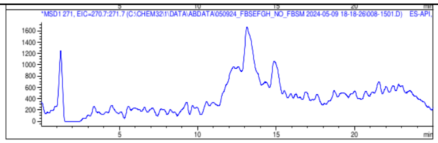 | 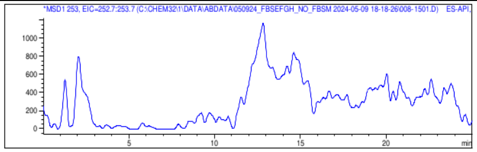 | 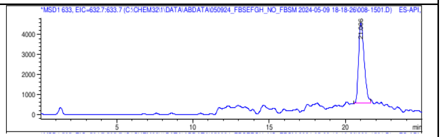 |
| | V348A 1 | 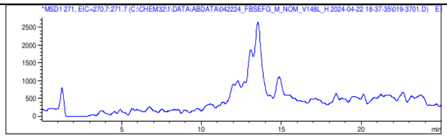 | 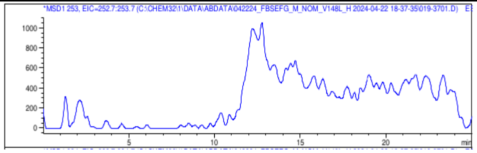 | 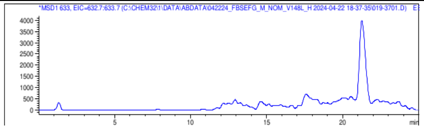 |
| | V348A 2 | 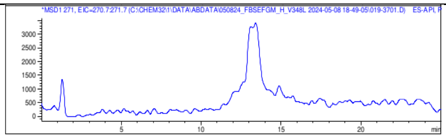 | 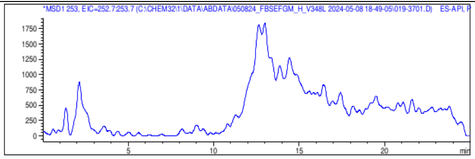 | 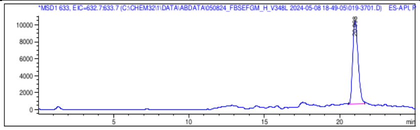 |
| | F350A 1 | 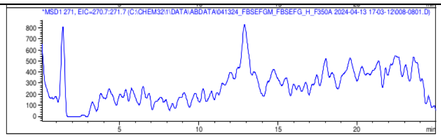 | 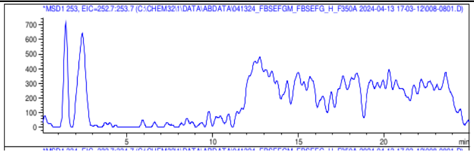 | 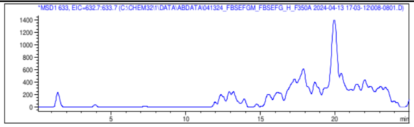 |
| | F350A 2 | 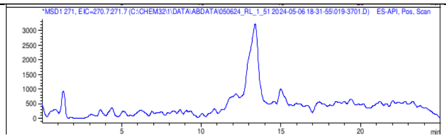 | 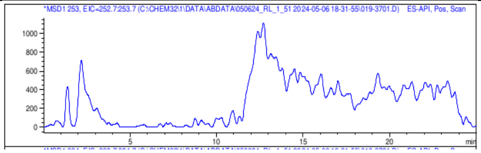 | 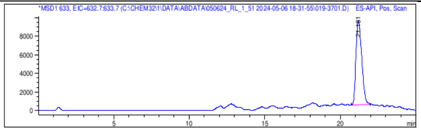 |
| | Standard | 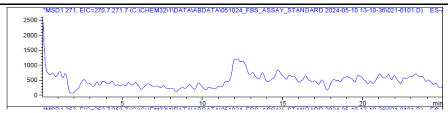 | 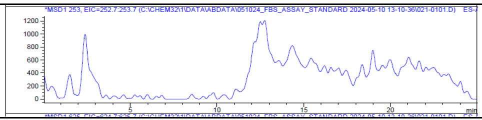 | 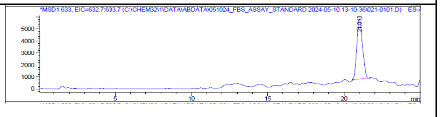 |

Supplemental Figure 6i. 4-Azidosalicylic acid

| | | Aryl-Ser | Aryl-Oxa | Fimsbactin A |
|---|---|---|---|---|
| 4-Azido Sal | FbsH Wildtype 1 | 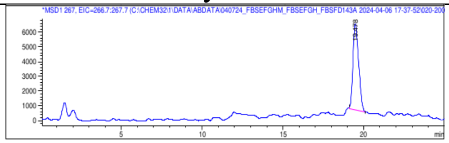 | 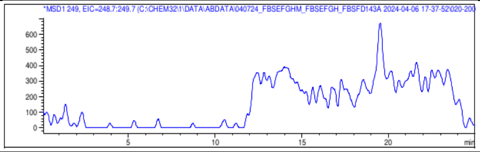 | 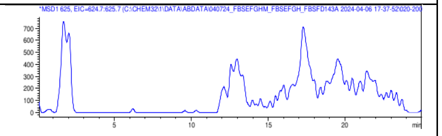 |
| | FbsH Wildtype 2 | 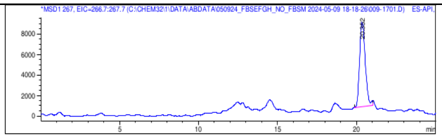 | 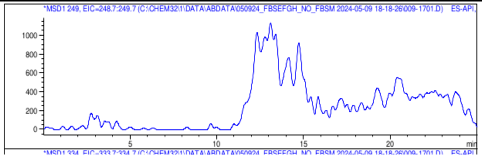 | 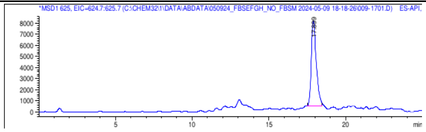 |
| | V348A 1 | 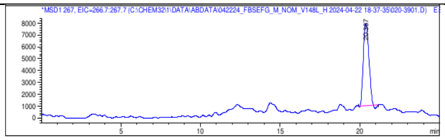 | 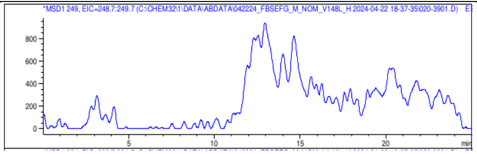 | 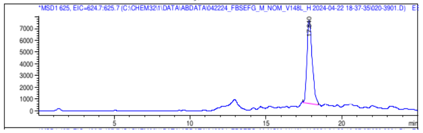 |
| | V348A 2 | 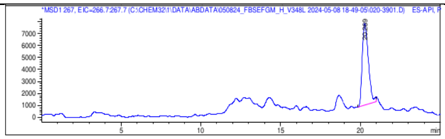 | 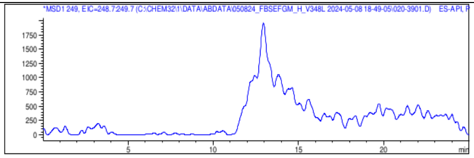 | 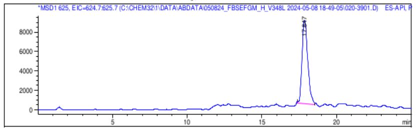 |
| | F350A 1 | 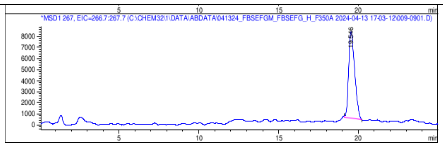 | 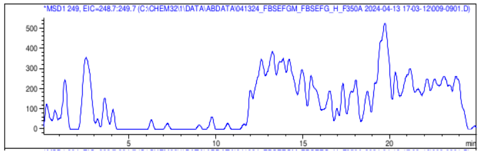 | 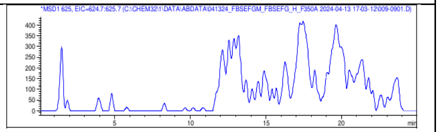 |
| | F350A 2 | 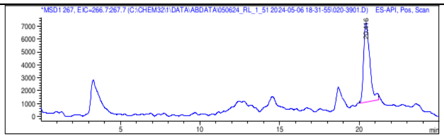 | 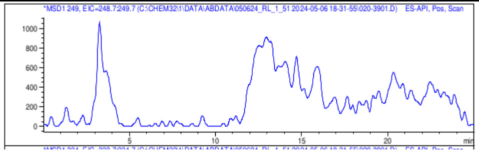 | 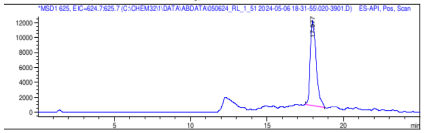 |
| | Standard | 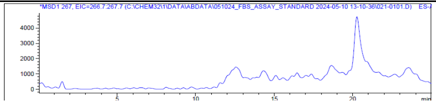 | 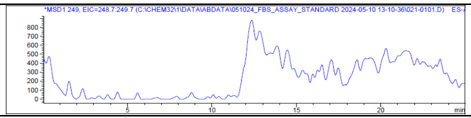 | 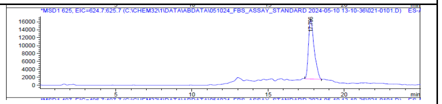 |

Supplemental Figure 6j. 4-Aminosalicylic acid

| | | Aryl-Ser | Aryl-Oxa | Fimsbactin A |
|---|---|---|---|---|
| 4-Amino Sal | FbsH Wildtype 1 | 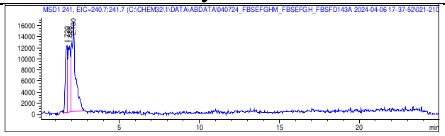 | 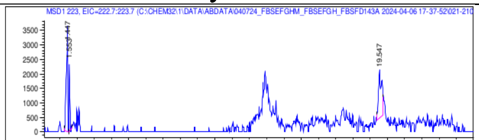 | 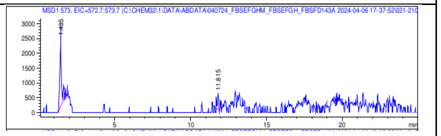 |
| | FbsH Wildtype 2 | 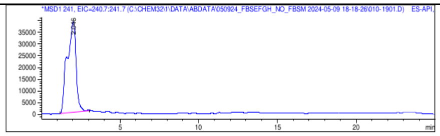 | 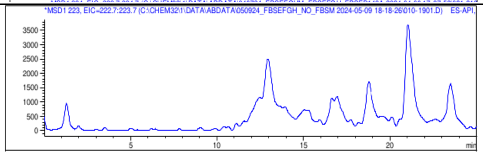 | 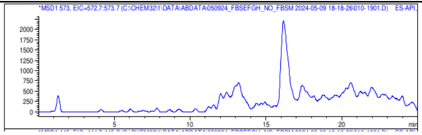 |
| | V348A 1 | 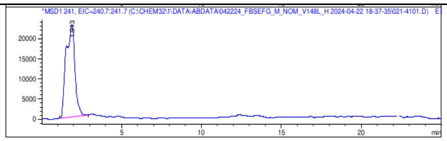 | 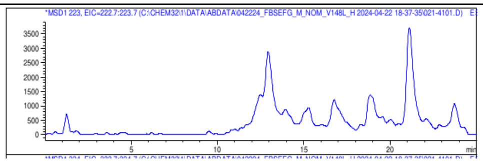 | 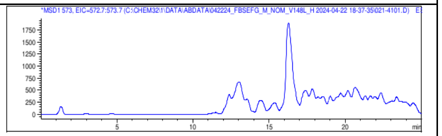 |
| | V348A 2 | 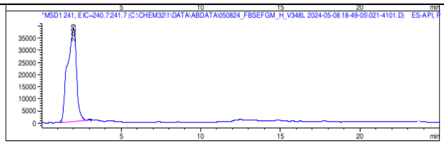 | 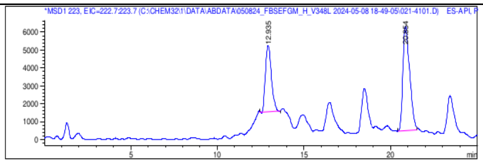 | 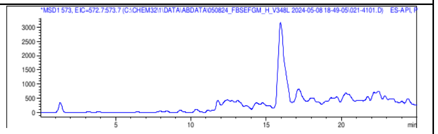 |
| | F350A 1 | 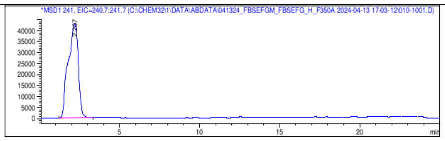 | 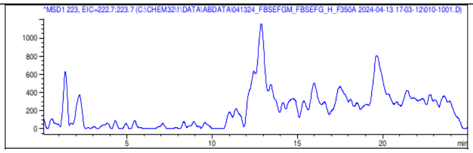 | 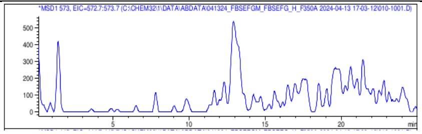 |
| | F350A 2 | 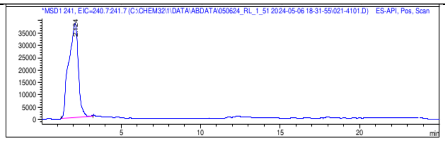 | 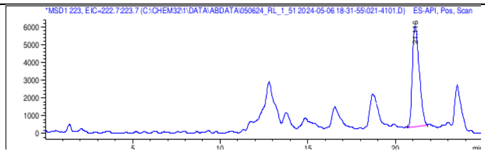 | 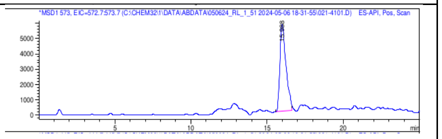 |
| | Standard | 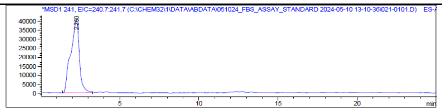 | 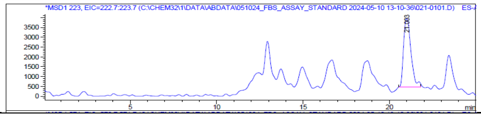 | 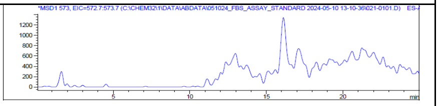 |

Supplemental Figure 6k. 4-Methylsalicylic acid

| | | Aryl-Ser | Aryl-Oxa | Fimsbactin A |
|---|---|---|---|---|
| 4-Methyl Sal | FbsH Wildtype 1 | 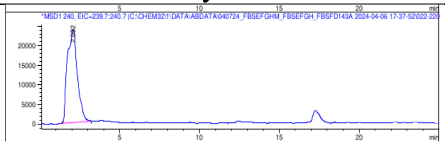 | 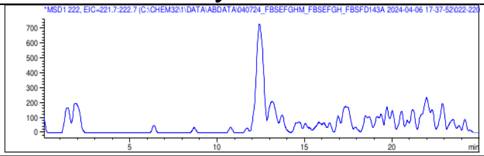 | 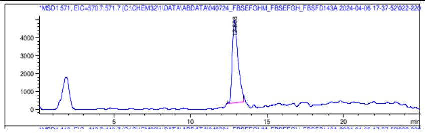 |
| | FbsH Wildtype 2 | 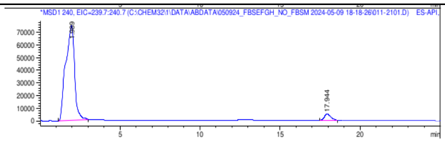 | 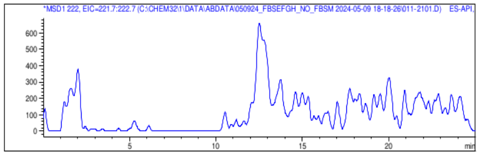 | 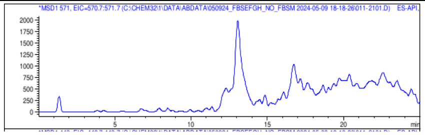 |
| | V348A 1 | 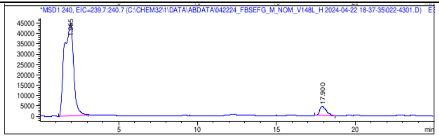 | 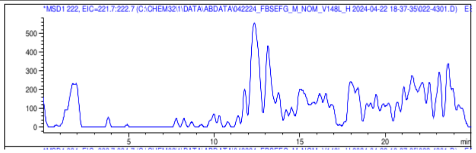 | 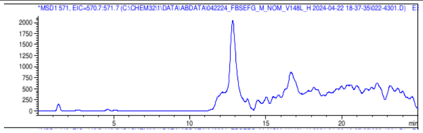 |
| | V348A 2 | 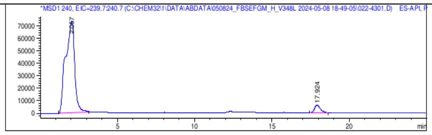 | 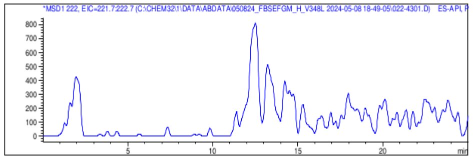 | 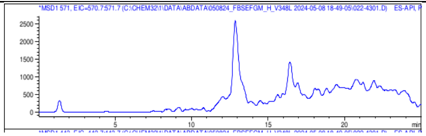 |
| | F350A 1 | 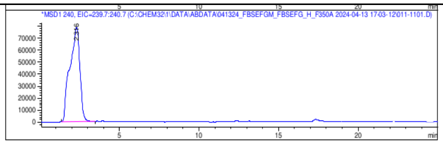 | 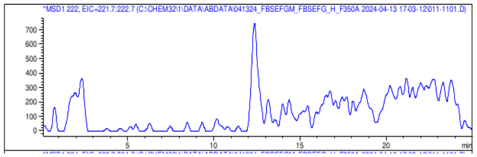 | 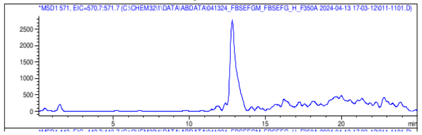 |
| | F350A 2 | 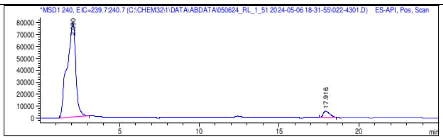 | 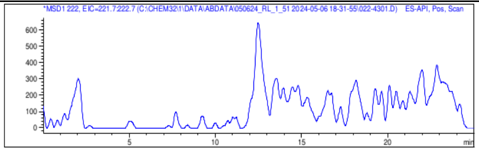 | 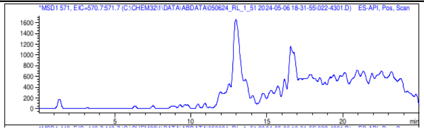 |
| | Standard | 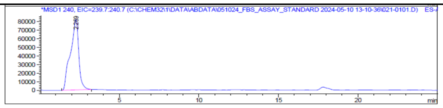 | 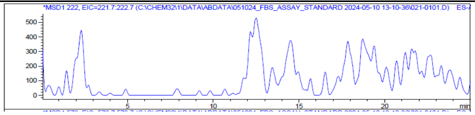 | 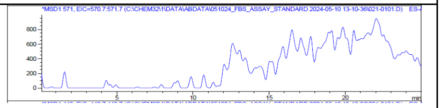 |

### Supplemental Tables

**Supplemental Table 1.** Sequence and structural similarity of FbsH with other structurally-characterized adenylation domains

| Protein | PDB | Conformation | FbsH | EntE | BasE | DhbE | MbtA | PchD | CahJ | NpsA | PqsA |
|---|---|---|---|---|---|---|---|---|---|---|---|
| **FbsH** | **9BHZ** | Intermediate | | 39.6% | 37.6% | 42.3% | 38.4% | 37.7% | 39.7% | 20.4% | 24.5% |
| **EntE** | **3RG2** | Thioester-forming | 0.84 Å | | 54.3% | 46.0% | 41.1% | 44.8% | 45.9% | 24.8% | 24.5% |
| **BasE** | **3O82** | C-term Disordered | 0.89 Å | 0.59 Å | | 42.5% | 40.0% | 40.2% | 42.1% | 22.8% | 21.2% |
| **DhbE** | **1MD9** | Adenylation | 0.86 Å | 0.82 Å | 0.82 Å | | 47.3% | 50.2% | 58.1% | 23.0% | 23.5% |
| **MbtA** | **5KEI** | Intermediate | 0.79 Å | 0.90 Å | 0.88 Å | 0.96 Å | | 45.0% | 57.7% | 24.7% | 28.6% |
| **PchD** | **7TYB** | Adenylate-forming | 0.83 Å | 0.70 Å | 0.70 Å | 0.81 Å | 0.73 Å | | 55.8% | 25.4% | 27.3% |
| **CahJ** | **5WM2** | Thioester-forming | 0.82 Å | 0.76 Å | 0.74 Å | 0.76 Å | 0.52 Å | 0.53 Å | | 26.6% | 25.1% |
| **NpsA** | **6VHT** | C-term Missing | 2.01 Å | 1.74 Å | 2.17 Å | 2.04 Å | 2.13 Å | 2.09 Å | 2.09 Å | | 21.5% |
| **PqsA** | **5OE3** | C-term Missing | 1.92 Å | 1.74 Å | 1.75 Å | 1.85 Å | 1.76 Å | 1.94 Å | 1.80 Å | 1.91 Å | |

**Supplemental Table 2**. Data collection and refinement statistics

| DATA COLLECTION | FbsH, DHB | FbsH, Sal-AMS |
|---|---|---|
| PDB CODE | **9BHY** | **9BHZ** |
| Beamline | APS 23-ID-D | APS 23-ID-D |
| Wavelength (Å) | 1.03373 | 1.03373 |
| Resolution range (Å) | 35.6 - 2.2 (2.3 - 2.2) | 35.5 - 2.2 (2.3 - 2.2) |
| Space group | P 1 | P 1 |
| a, b, c (Å) | 56.35 74.61 75.46 | 56.33 74.17 75.56 |
| α, β, γ (°) | 85.03 69.87 70.28 | 85.05 69.91 70.09 |
| Total reflections | 187670 (17902) | 181406 (17705) |
| Unique reflections | 53810 (5285) | 52017 (5132) |
| Multiplicity | 3.5 (3.4) | 3.5 (3.4) |
| Completeness (%) | 97.92 (95.02) | 97.94 (96.05) |
| Mean I/sigma(I) | 9.24 (1.11) | 9.90 (1.19) |
| $R_{merge}$ | 0.1411 (1.197) | 0.1871 (1.183) |
| $R_{pim}$ | 0.08961 (0.7697) | 0.1201 (0.7654) |
| $CC_{1/2}$ | 0.989 (0.231) | 0.968 (0.714) |
| **Refinement** | | |
| Resolution range (Å) | 35.6 - 2.2 (2.3 - 2.2) | 35.5 - 2.2 (2.3 - 2.2) |
| Reflections, refinement | 53693 (5209) | 52019 (5132) |
| Reflections, $R_{free}$ | 1990 (185) | 1995 (204) |
| $R_{work}$ | 0.1942 (0.2758) | 0.1974 (0.2798) |
| $R_{free}$ | 0.2310 (0.3263) | 0.2388 (0.3317) |
| Protein residues | 1031 | 986 |
| Ligands of interest | 2 DHB | 2 Sal-AMS |
| Other molecules | 2 EDO | 2 EDO |
| Water molecules | 142 | 182 |
| Rms (bonds) (Å) | 0.019 | 0.008 |
| Rms (angles) (°) | 0.81 | 0.87 |
| Rama. Favored (%) | 97.73 | 97.15 |
| Rama. Allowed (%) | 2.27 | 2.85 |
| Rama. Outliers (%) | 0.00 | 0.00 |
| Rotamer outliers (%) | 0.76 | 1.87 |
| Average B-factor ($Å^2$) | | |
| overall, protein | 47.99 | 50.32 |

**Supplemental Table 3:** Primers used in the study

| *Primer name* | **Sequence 5'-3'** |
|---|---|
| *FbsH_V348A_F* | GGCTGAAGGTTTGGCGAATTTCACACACTTGG |
| *FbsH_V348A_R* | CCAAGTGTGTGAAATTCGCCAAACCTTCAGCC |
| *FbsH_F350A_F* | GGTGAATGCCACACACTTGGATGATTCAG |
| *FbsH_F350A_R* | CTGAATCATCCAAGTGTGTGGCATTCACC |

**Supplemental table 4.** Codon-optimized gene encoding FbsH.

| Genes | Sequence |
|---|---|
| *fbsH* | ATGTTTATAGAACAAGATATTCAACTGGGCTATGTGCCATTTCCCAAATCGAGAGCTCAGCAGTATCGA<br>GATGAAGGTTGCTGGAAATCTCAAACCCATTTTCAATTGTTGGCAAGTTTAAAAGACCGCTTTGCACAG<br>CGTGTAGCCGTCATACAAGATGATAAGCAACTGACCTATCAGCAGTTATATGACTATGCGATTCATTAT<br>GGCACTTACTTAAAACAACAGGGTATTCGTGAAACCGACTTTGTTTTATTGCAGTCGCCGAATGTGATT<br>GAAGTTTTTATTGTGATTTTTGGTTTATATGCCATTGGTGCAAGACCTGTATTTTGCTTGCATGGTCAT<br>GGCTCATATGAAATTGAAAATATTGCAAGACAAAGCCGTGCAGTAGGTTTTCTTAAACTGTGTGGTTCC<br>GCCAATGAAAGTACTGCAACCGACGTTTGTGAAGAATTTTCTAAACCCAATTTTAAGTTGTGGTTTAGA<br>GAAAGTATTGTGAGTCGATCGAGCATTGAAGCGTCATTGCCACAGTTACAAGGTGTTGCGCCTGCCTTC<br>AATTTACGCGCACAGAGTGAATCTGAAGATATCGCATTTTTACAACTTTCAGGTGGAACCACAGGTTTA<br>CCAAAACTGATTCCTCGCACTCATGCGGACTACATTTACTCGATTGAAAAAAGCGTCGACGTTGCTGGG<br>CTAACTCAGGACACCAAACAGTTGGTCGTACTACCTGTGATGCATAATTTCTGCATGAGCTCACCTGGG<br>TTTCTGGGGGTGTTTTATGTAGGTGGCACAGTGGTACTCAGTCAGTTGACACATCCTCGTGTTTGTTTT<br>GAATTGATTGAAAAATACCAAATTCAGCAAGTCTCTTTAGTTCCTGCTATTGCCACGCTTTGGCTCAAT<br>GCTGAAAGTCTCAAAGATTATGATTTATCCAGCTTGCAAGTGGTGCAAGTCGGTGGTGCCAAATTGTTA<br>CCTTCGCTTGCCGAGCAGATTATTGACACCCTGCAAGTCAAATTGCAGCAAGTCTATGGCATGGCTGAA<br>GGTTTGGTGAATTTCACACACTTGGATGATTCAGACCAAATTACCATTCAAACCCAAGGTAAAAAGCTG<br>TCGCATTTAGATGAAATCCGTATTGCTGATCAGGATGGCAATGCTTTGCCAATCAATGCGATCGGGCAT<br>ATTCAAACCCGTGGTCCTTATACCATCAATGGGTATTACAACTTGCCTGAAATTAACCAGCGTGCCTTT<br>ACGCAAGATGGCTTCTATAAAACGGGTGATATTGGTTATTTAGATGAAAATCTGAATATTGTGGTTACG<br>GGGCGTGAAAAGGAACAGATCAACCGTTCTGGTGAAAAAATCACACCAAGTGAAATTGAAGAATTTATT<br>TTGCAATATCCATCGGTGAAAGATGTTTGTGTGATTGGCGTCAGTGACGATTATTTAGGTGAACGTATT<br>AAAGCCATCATCATTCCAAAGTTAGATGACAGTGAAATCAACTTAAAAAATATTCGTAAATTTTTGATT<br>AGTAAAAATATTGCGCATTTCAAAATCCCTGATGAAATCGAAGTCGTGGCAGATTTTAAATATACCCAC<br>GTGGGCAAAGTCAATCGACAAAAACTGGGATAA |

**Supplemental table 5.** Protein sequence

| Protein | Sequence |
|---|---|
| *FbsH (WP_049594809.1)* | MGSSHHHHHHSSGLVPRGSHMASMTGGQQMGRGSIMFIEQDIQLGYVPFPKSRAQQYRDEGCWKSQ<br>THFQLLASLKDRFAQRVAVIQDDKQLTYQQLYDYAIHYGTYLKQQGIRETDFVLLQSPNVIEVFIV<br>IFGLYAIGARPVFCLHGHGSYEIENIARQSRAVGFLKLCGSANESTATDVCEEFSKPNFKLWFRES<br>IVSRSSIEASLPQLQGVAPAFNLRAQSESEDIAFLQLSGGTTGLPKLIPRTHADYIYSIEKSVDVA<br>GLTQDTKQLVVLPVMHNFCMSSPGFLGVFYVGGTVVLSQLTHPRVCFELIEKYQIQQVSLVPAIAT<br>LWLNAESLKDYDLSSLQVVQVGGAKLLPSLAEQIIDTLQVKLQQVYGMAEGLVNFTHLDDSDQITI<br>QTQGKKLSHLDEIRIADQDGNALPINAIGHIQTRGPYTINGYYNLPEINQRAFTQDGFYKTGDIGY<br>LDENLNIVVTGREKEQINRSGEKITPSEIEEFILQYPSVKDVCVIGVSDDYLGERIKAIIIPKLDD<br>SEINLKNIRKFLISKNIAHFKIPDEIEVVADFKYTHVGKVNRQKLG |

The sequence highlighted in red represents N-His$_6$-tag and the thrombin cleavage site residues (LVPR^GS) contributed by vector pET28a.
